# Opposing actions of JIP4 and RILPL1 provide antagonistic motor force to dynamically regulate membrane reformation during lysosomal tubulation/sorting driven by LRRK2

**DOI:** 10.1101/2024.04.02.587808

**Authors:** Luis Bonet-Ponce, Tsion Tegicho, Alexandra Beilina, Jillian H. Kluss, Yan Li, Mark R. Cookson

## Abstract

Lysosomes are dynamic cellular structures that adaptively remodel their membrane in response to stimuli, including membrane damage. We previously uncovered a process we term LYTL (LYsosomal Tubulation/sorting driven by Leucine-Rich Repeat Kinase 2 [LRRK2]), wherein damaged lysosomes generate tubules sorted into mobile vesicles. LYTL is orchestrated by the Parkinson’s disease-associated kinase LRRK2 that recruits the motor adaptor protein and RHD family member JIP4 to lysosomes via phosphorylated RAB proteins. To identify new players involved in LYTL, we performed unbiased proteomics on isolated lysosomes after LRRK2 kinase inhibition. Our results demonstrate that there is recruitment of RILPL1 to ruptured lysosomes via LRRK2 activity to promote phosphorylation of RAB proteins at the lysosomal surface. RILPL1, which is also a member of the RHD family, enhances the clustering of LRRK2-positive lysosomes in the perinuclear area and causes retraction of LYTL tubules, in contrast to JIP4 which promotes LYTL tubule extension. Mechanistically, RILPL1 binds to p150^Glued^, a dynactin subunit, facilitating the transport of lysosomes and tubules to the minus end of microtubules. Further characterization of the tubulation process revealed that LYTL tubules move along tyrosinated microtubules, with tubulin tyrosination proving essential for tubule elongation. In summary, our findings emphasize the dynamic regulation of LYTL tubules by two distinct RHD proteins and pRAB effectors, serving as opposing motor adaptor proteins: JIP4, promoting tubulation via kinesin, and RILPL1, facilitating tubule retraction through dynein/dynactin. We infer that the two opposing processes generate a metastable lysosomal membrane deformation that facilitates dynamic tubulation events.

## INTRODUCTION

Lysosomes were initially discovered by Christian de Duve (de Duve et al., 1955) as acidic organelles responsible for the hydrolysis of macromolecules. Beyond cargo clearance, lysosomes play a multifaceted role within the cell (Ballabio and Bonifacino, 2020). Lysosomes control nutrient responses (Goul et al., 2023), regulate the morphology of other organelles (Spits et al., 2021; Wong et al., 2018), facilitate cholesterol egress (Chu et al., 2021; Höglinger et al., 2019), and co-transport other cellular structures (Liao et al., 2019; Guo et al., 2018). Thus, lysosomes serve as central organelles that help maintain cellular homeostasis, particularly during periods of cellular stress (Tan and Finkel, 2023).

When the integrity of the lysosomal membrane is compromised, cells can initiate "lysosomal cell death" (Wang et al., 2018). This process involves the leakage of protons and hydrolases into the cytosol, ultimately damaging multiple cellular macromolecules and terminating cell viability. We and others have characterized different pathways by which lysosomes deal with membrane damage. When the damage is limited, lysosomes can repair their membrane through ESCRT (Skowyra et al., 2018; Radulovic et al., 2018; Herbst et al., 2020) or lipid exchange with the endoplasmic reticulum (Tan and Finkel, 2022; Radulovic et al., 2022). RNA granules can also plug rupture sites to prevent lysosomes from collapsing and enable an efficient repair response (Bussi et al., 2023). When damage persists, cells clear ruptured lysosomes via selective autophagy in a process known as lysophagy (Maejima et al., 2013). Additionally, lysosomal membrane damage can induce a non-canonical autophagy pathway called CASM (conjugation of ATG8 to single membranes) (Kaur et al., 2023; Boyle et al., 2023; Corkery et al., 2023), which might be associated with lysosomal membrane repair (Cross et al., 2023). Collectively, the existence of these multiple, partially redundant pathways indicate how critical it is for cells to avoid lysosomal membrane damage.

Beyond membrane repair and autophagy, lysosomal membrane damage triggers a unique tubulation and membrane sorting process, which we have named LYTL (LYsosomal Tubulation/sorting driven by LRRK2) (Bonet-Ponce et al., 2020; Bonet-Ponce and Cookson, 2022b). LYTL is orchestrated by Leucine-rich repeat kinase 2 (LRRK2), a large kinase basally located in the cytosol that is activated at membranes (Bonet-Ponce and Cookson, 2022a) where it phosphorylates a subset of RAB GTPases (Steger et al., 2016; Kluss et al., 2022b). Recent evidence indicates that LRRK2 is recruited to ruptured lysosomes via RAB12 (Wang et al., 2023; Dhekne et al., 2023) and the CASM machinery (Bentley-DeSousa and Ferguson, 2023; Eguchi et al., 2024), where it phosphorylates and brings pRAB proteins to the lysosomal membrane (Eguchi et al., 2018; Herbst et al., 2020; Bonet-Ponce et al., 2020). Subsequently, pRAB10 recruits its effector JIP4 (C-jun-amino-terminal kinase-interacting protein 4, a motor adaptor protein) that promotes the formation of a lysosomal tubule negative for lysosomal membrane markers (Bonet-Ponce et al., 2020; Bonet-Ponce and Cookson, 2022b; Kluss et al., 2022a). The JIP4-positive LYTL tubule can be resolved into vesicles that travel through the cytosol and contact healthy lysosomes. LYTL is regulated by lysosomal positioning (Kluss et al., 2022a), and the membrane sorting step is controlled by the endoplasmic reticulum (Bonet-Ponce and Cookson, 2022b). However, the entirety of molecular pathways that regulate LYTL remain undefined.

Here, we use unbiased proteomics on isolated lysosomes to characterize the repertoire of proteins whose abundance on lysosomes depend on the activity of LRRK2. We show that LRRK2 recruits RILP-like protein 1 (RILPL1) to ruptured lysosomes in a pRAB-dependent manner. RILPL1 is a RILP-Homology Domain (RHD) family member that clusters LRRK2-positive lysosomes to the perinuclear area and retracts LYTL tubules. Mechanistically, RILPL1 binds to p150^Glued^ (a dynactin subunit), enhancing its recruitment to the damaged lysosomes, and promoting the transport of lysosomes and tubules to the minus-end of microtubules. By further characterizing the tubulation process, we demonstrate that LYTL tubules move along tyrosinated microtubules and tubulin tyrosination is required for tubule elongation. In summary, our findings underscore the dynamic regulation of LYTL tubules by two distinct pRAB effectors, functioning as opposing motor adaptor proteins: JIP4, which promotes tubulation, and RILPL1, which facilitates tubule retraction. Collectively, these two opposing forces lead to a metastable membrane structure that supports a dynamic and transient tubulation event.

## RESULTS

### Using unbiased proteomics to uncover the LRRK2 lysosomal proteome

To establish a comprehensive map of the LRRK2 lysosomal proteome, we created a HEK293T cell line that stably expresses GFP-LRRK2 and TMEM192-3xHA. TMEM192 is a lysosomal membrane protein used as a bait to immuno isolate lysosomes in a technique known as LYSO-IP (Abu-Remaileh et al., 2017). We confirmed that, in the presence of the lysosomotropic damaging agent LLOME (L-leucyl-L-leucine methyl ester), LRRK2 colocalized with TMEM192 (Fig. S1A-B). We also confirmed the presence of endogenous RAB10 and JIP4 on lysosomes, which were completely abrogated by LRRK2 kinase inhibition using MLi2 (Fig. S1C-D). We next validated the LYSO-IP as a suitable technique to isolate lysosomes (Fig. S1E-G) and to capture the enrichment of pRAB10, total RAB10 and JIP4 to the lysosomal fraction in a LRRK2 kinase-dependent manner (Fig. S1F).

We reasoned that any proteins recruited by LRRK2 to lysosomes would be present on isolated lysosomes from LLOME-treated cells, but absent in cells pre-treated with MLi2 (Fig. 1A-B). To validate that we could detect changes in lysosomal proteome, we compared DMSO- and LLOME-treated cells. As expected, LLOME addition promoted the recruitment of members of the ESCRT complex, known autophagy/lysophagy proteins (Eapen et al., 2021), and LRRK2. We also observed a significant decrease in lysosomal hydrolases upon LLOME treatment (Fig. S1I-J). Notably LRRK2 kinase inhibition did not modify the lysosomal presence of any of the ESCRT members, autophagy proteins or lysosomal hydrolases analyzed (Fig. S1I-J) suggesting that LRRK2 is not involved in maintaining the integrity of the lysosomal membrane nor promoting lysosomal exocytosis. We also found that 46 proteins were significantly increased on lysosomes from cells treated with LLOME compared to co-treatment with LLOME and MLi2 (Fig. 1C). The 46 candidates were enriched for membrane trafficking terms (Fig. 1D), including “vesicle docking involved in exocytosis”, “protein secretion”, "response to calcium”, “protein localization to plasma membrane” and “establishment of vesicle localization”. Specifically, nine RAB GTPases were identified in our screen (RAB3A/B/D, RAB8A/B, RAB10, RAB12, RAB29 and RAB35) (Fig. 1C,E,F; Fig. S2A). All of these RABs have been proposed as LRRK2 substrates (Steger et al., 2016) and no other RAB protein was identified. Interestingly, RAB5A/B/C and RAB43 (Fig. 1F; Fig. S2A) were also originally proposed as LRRK2 substrates but were not recovered in our survey suggesting that these RABs are not *bona fide* substrates in the context of lysosomal damage. We also identified eight calcium-binding proteins, six RNA-binding proteins and eight dynein-associated proteins (Fig. 1E). Consistent with our previous work, we detected JIP4 as a positive hit, further validating our screening strategy (Fig. 1C,E,F; Fig. S2B). RILPL1 was detected as a new LRRK2- interacting candidate on lysosomes (Fig. 1C,E,F; Fig. S1K; Fig. S2B). RILPL1 is a RILP-homology domain (RHD) family member, along with RILPL2, JIP3 and JIP4. All four proteins are predicted to act as pRAB effectors (Waschbüsch et al., 2020). RILPL1 has been previously linked to LRRK2 and has been reported to reduce ciliogenesis and alter centrosomal cohesion (Dhekne et al., 2018; Lara Ordónez et al., 2019). Therefore, we further examined RILPL1 as a potential mediator of responses to lysosomal membrane damage.

**Figure 1.**
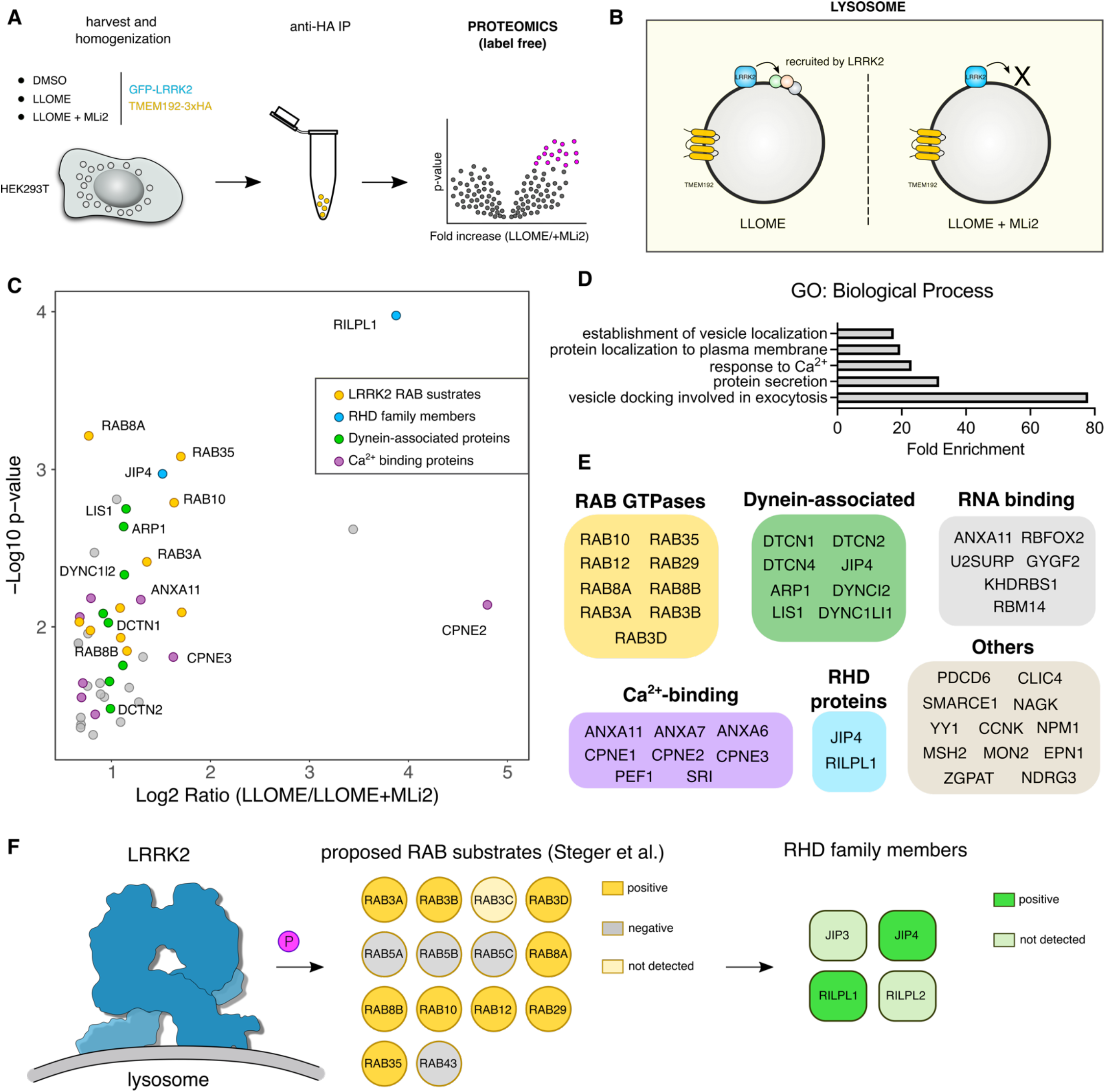
Unbiased characterization of the LRRK2 lysosomal proteome. (A-B) Schematic depiction of the experimental workflow of the lysosomal immunoprecipitation (LYSO-IP) in HEK293T stably expressing LRRK2, reasoning that the proteins recruited by LRRK2 to damaged lysosomes will be absent if the cells are pre-treated with the LRRK2 kinase inhibitor MLi2. (C) Volcano plot showing the subset of proteins with a Fold Change > 1.3 and p value < 0.05 present on the lysosomal fraction of cells treated with LLOME vs LLOME+MLi-2. Data are from 4 independent experiments. (D) Gene Ontology (GO) search of the top 5 enriched terms for Biological Process of the 46 candidates found in (C), with Fold Enrichment indicated on the horizontal axis. (E) The 46 candidate proteins were grouped by molecular function. (F) Model of the LRRK2 lysosomal proteome, where LRRK2 recruits nine RAB substrates and two RHD family members to damaged lysosomes.

### LRRK2 recruits RILPL1 to lysosomes via pRAB proteins

We used immunostaining as an orthogonal approach to validate that the recruitment of RILPL1 to lysosomes occurs across cell types in a manner that depends on LRRK2. Endogenous RILPL1 was detected on LRRK2- positive lysosomes after LLOME treatment in HEK293T cells, U2OS cells and mouse primary astrocytes (Fig. S3A-B; Fig. 2A). As expected, we could not detect RILPL1 presence on lysosomes after LRRK2 kinase inhibition (Fig. S3A; Fig. 2A-B). The anti-RILPL1 antibody used in this study was validated against siRNA for western blot and immunocytochemistry (Fig. S3C-D). RILPL1 recruitment to LRRK2-positive lysosomes was also observed in live cells (Fig. 2C). The presence of active LRRK2 on lysosomes is sufficient to recruit RILPL1 as we observed RILPL1 presence on lysosomes when trapping LRRK2 to the lysosomal membrane using a previously described chimera construct expressing the first 39 amino acids of LAMTOR1 (LYSO-LRRK2 plasmid) (Kluss et al., 2022a) (Fig. S3E), thereby bypassing the need to damage lysosomal membranes using LLOME. We hypothesized that RILPL1 is recruited to LRRK2-positive lysosomes via pRAB proteins. Supporting this contention, we observed a physical interaction through immuno-precipitation between a tagged version of RILPL1 and pRAB10 in LLOME- treated cells (Fig. 2D). This interaction was further reproduced in U2OS cells (Fig. S3F) and was, again, dependent on LRRK2 kinase activity (Fig. S3G).

**Figure 2.**
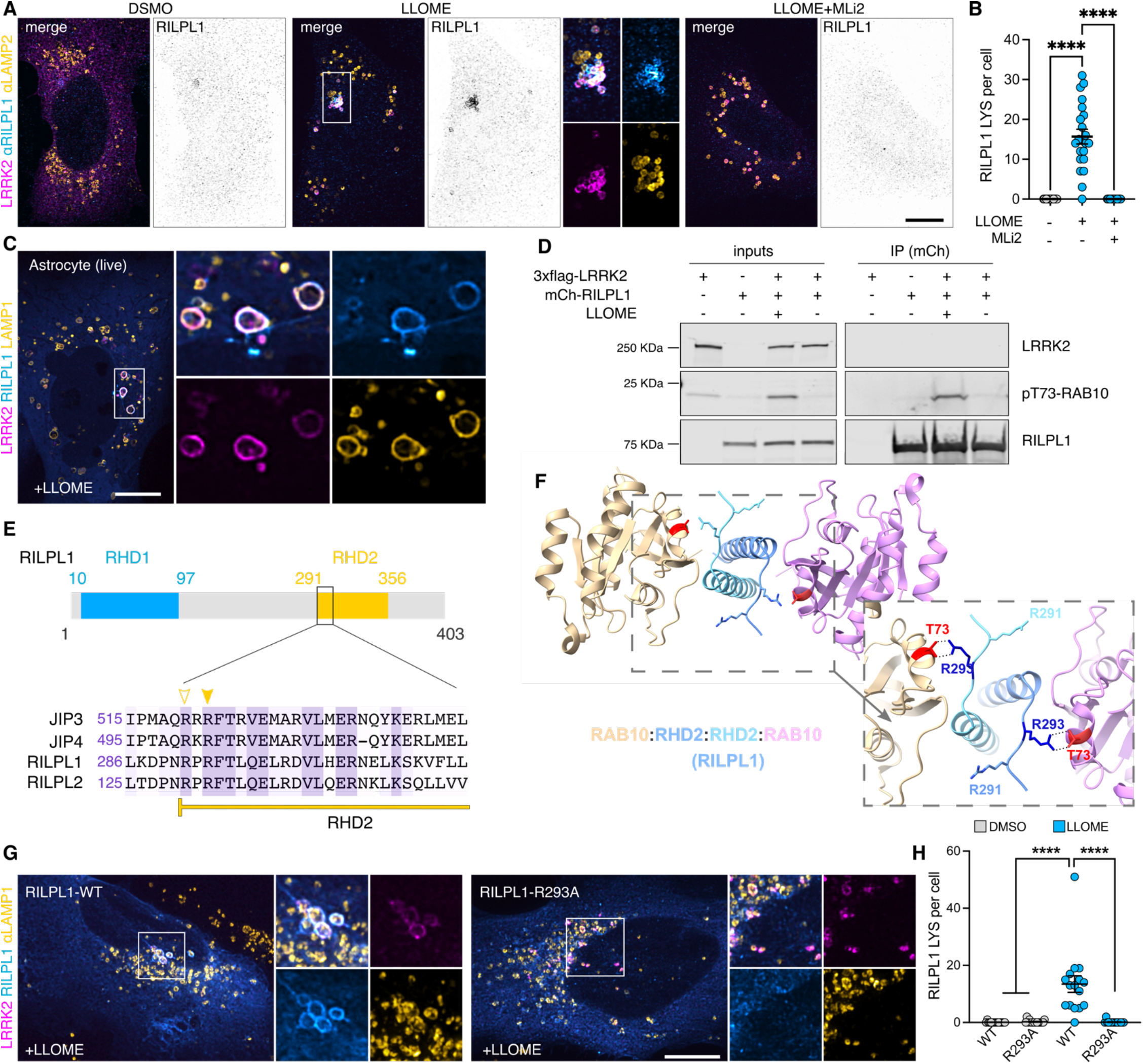
LRRK2 recruits RILPL1 to damaged lysosomes via pRAB proteins. (A) U2OS cells were transfected with HaloTag-LRRK2 for 36 h. Cells were then treated with DMSO, LLOME or LLOME+MLi2 for 2 h before fixation. Cells were then stained for RILPL1 and LAMP2. (B) Quantification of the RILPL1 lysosomes per cell in the three different conditions. Data are mean ± SEM (p<0.0001, n = 21 cells). One-way ANOVA with Tukey’s post hoc tests. (C) Mouse primary astrocytes were transfected with HaloTag-LRRK2, mCherry- RILPL1 and LAMP1-mNeonGreen for 48 h. Astrocytes were treated with LLOME for 6 h and imaged live. (D) HEK293FT cells were transfected with 3xflag-LRRK2 and mCherry-RILPL1 for 24 h. Cells were treated or not with LLOME for 2 h and lysates were subjected to immunoprecipitation with anti-RFP antibodies. (E) RILP-Homoloy Domain 2 (RHD2) alignment of JIP3, JIP4, RILPL1 and RILPL2 using Clustal (Clustal Omega, EMBL). Arrowheads show the conserved Arg. residues responsibles for the binding with the pRAB substrates. Filled arrowhead marks the Arg. that binds with the phosphorylated residue, and the empty arrowhead indicates the Arg. that stabilizes the interaction. (F) Structural model of the interaction between the RHD2 of RILPL1 and RAB10, showing the interaction between T73- RAB10 and R293-RILPL1. (G) U2OS cells were transfected with HaloTag-LRRK2 and mCherry-RILPL1 (WT or R293A) for 36 h. Cells were then treated with DMSO or LLOME for 2 h, fixed and stained for LAMP1. (H) Quantification of RILPL1-positive lysosomes per cell in the different groups. Data are mean ± SEM (p<0.0001, n = 16 cells). One-way ANOVA with Tukey’s post hoc test. Scale bar (A,G)= 10 µm; (C)= 20 µm.

JIP3, JIP4, RILPL1, and RILPL2 have two conserved Arginine (R) residues in their RILP-homology domain 2 (RHD2) that are predicted to mediate the binding to the phosphorylated residue of the pRAB substrates (Waschbüsch et al., 2020) (Fig. 2E). Thus, this group of proteins is proposed to act as pRAB effectors. Previous *in vitro* structural biology studies have demonstrated that RILPL2 binds to RAB8A-pThr.72 through an exposed arginine (R132) in its RHD2 region (Waschbüsch et al., 2020). The binding is enhanced by a nearby Arg. residue (R130) (Waschbüsch et al., 2021). Given that both R residues are conserved in RILPL1, we considered that a similar binding mechanism might occur (Fig. 2E). Our structural model confirms that RILPL1 binds to pThr.73 through R293 forming a tetramer (Fig. 2F). When Arg.293 was mutated into alanine (RILPL1-R293A), LRRK2 was unable to recruit RILPL1 to the lysosomal membrane in cells treated with LLOME (Fig. 2G-H). Altogether, our data strongly suggest that RILPL1 is a pRAB effector, with similar mode of binding to the previously described effector JIP4.

### RILPL1 clusters LRRK2-positive lysosomes to the centrosome and reduces LYTL tubulation

As we previously reported, LRRK2-positive lysosomes cluster around the centrosome upon LLOME treatment (Kluss et al., 2022a) (Fig. 3A). Interestingly, in cells lacking RILPL1, the clustering of LRRK2-positive lysosomes around the centrosome was drastically reduced (Fig. 3B-C) even though they are still close to the nucleus (Fig. 3B,D). We next wondered if LRRK2 recruitment is restricted to lysosomes clustered around the centrosome, or alternatively if LRRK2 recruitment precedes and promotes perinuclear clustering. Time-lapse microscopy experiments demonstrated that LRRK2 recruitment proceeds the clustering of LRRK2-positive lysosomes in the perinuclear space (Fig. 3E, Suppl. Movie 1). In U2OS cells lacking exogenous LRRK2 expression, RILPL1 knockdown has no effect in lysosomal positioning in untreated cells or LLOME-treated cells (Fig. S3H-J), suggesting that the role of RILPL1 on lysosomal positioning is LRRK2-dependent. Collectively, these data suggest that RILPL1 is responsible for clustering LRRK2-positive lysosomes around the centrosome. We have previously shown that perinuclear clustering promotes the ability of LRRK2 to recruit JIP4 and initiate LYTL (Kluss et al., 2022a). While RILPL1 knockdown does not affect LRRK2 recruitment to lysosomes (Fig. 3F-G), it did significantly decrease the presence of JIP4 on LRRK2-positive lysosomes (Fig. 3F,H,I). Thus, RILPL1 and JIP4 likely bind pRAB proteins using similar motifs but in an antagonistic manner.

**Figure 3.**
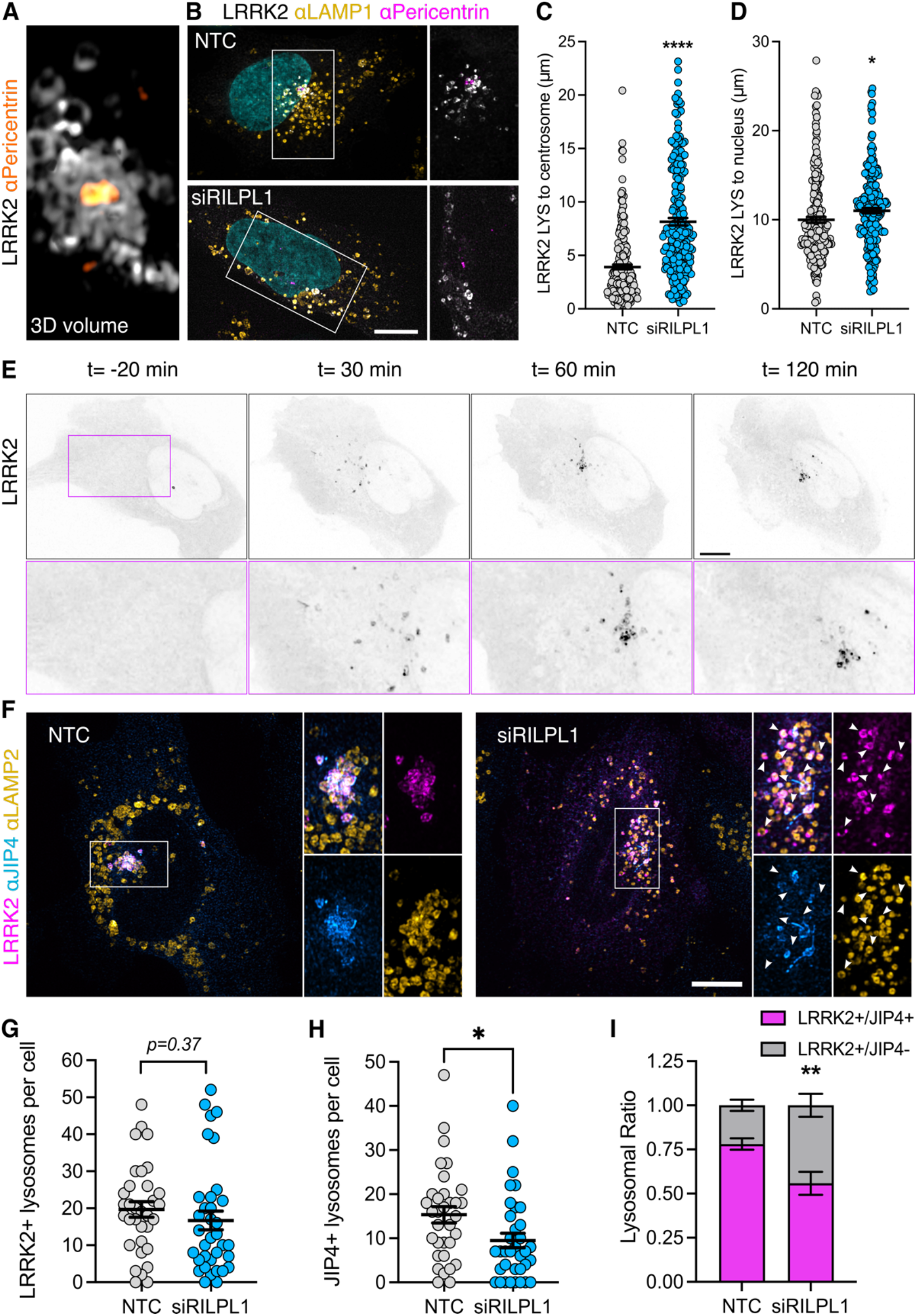
RILPL1 promotes LRRK2+ lysosome clustering around the centrosome. (A) U2OS cells were transfected with HaloTag-LRRK2 for 36 h. Cells were then treated with LLOME (2 h), fixed and stained with pericentrin. Volume view image of LRRK2+ lysosomes clustered around the centrosome (pericentrin). (B) U2OS cells were treated with a non-targeting control (NTC) or siRILPL1 RNA. 24 h later, cells were transfected with HaloTag-LRRK2 for 36 h. Then, cells were treated with LLOME (2 h), and were fixed and stained with a pericentrin antibody. (C) The distance between LRRK2+ lysosomes and the centrosome was measured in cells treated with NTC or siRILPL1. Data are mean ± SEM (p<0.0001, n = 221-222 lysosomes, from 7 and 9 cells). Unpaired t-test with Welch’s correction. (D) The distance between LRRK2+ lysosomes and the center of the nucleus was measured in cells treated with NTC or siRILPL1. Data are mean ± SEM (p= 0.0206, n = 221-222 lysosomes, from 7 and 9 cells). Unpaired t-test. (E) Time-lapse of a U2OS cell expressing HaloTag- LRRK2 and treated with LLOME. LRRK2 puncta recruitment and movement is followed in different time points up to 120 min. (F) U2OS cells were treated with NTC or siRILPL1 RNA. 24 h later, cells were transfected with HaloTag- LRRK2 for 36 h. Then, cells were treated with LLOME (2 h), and were fixed and stained with JIP4 and LAMP2 antibodies. (G) LRRK2+ lysosomes per cell were quantified. Data are mean ± SEM (n = 33-35 cells). Unpaired t-test. (H) JIP4+ lysosomes per cell were quantified. Data are mean ± SEM (p= 0.0202, n = 33-35 cells). Unpaired t-test. (I) The ratio of LRRK2+/JIP4+ lysosomes compared to LRRK2+/JIP4- lysosomes in NTC vs siRILPL1 treated cells. Data are mean ± SEM (p= 0.0035, n = 31-33 cells). Unpaired t-test with Welch’s correction for unequal variance. Arrowheads indicate LRRK2+/JIP4- lysosomes. Scale bar= 10 µm.

Based on these results, we considered if RILPL1 might also play a role in LYTL. Super-resolution microscopy identified RILPL1 tubules that emanate from LRRK2-positive lysosomes that were negative for LRRK2 and LAMP1 (Fig. 4A). Additionally, RILPL1 colocalizes with JIP4 on the lysosomal tubules demonstrating that both effectors bind the same LYTL-derived structures (Fig. 4B). The presence of RILPL1 on LYTL tubules was corroborated at the endogenous level (Fig. S3K), and is therefore not an artifactual consequence of overexpression. RILPL1 knockdown leads to a modest increase in tubule length (Fig. 4C-D) but a strong increase in tubular ratio (Fig.4C,E), which measures the number of JIP4-positive tubules divided by the number of JIP4- positive lysosomes in each cell. These results oppose the effect previously reported with JIP4 (Bonet-Ponce et al., 2020). We therefore infer that while both effector proteins bind pRABs at the LYTL tubules, JIP4 promotes tubule elongation and RILPL1 triggers tubule retraction, generating a dynamic metastable structure from the lysosomal surface. Consistently, RILPL1 depletion leads to a decrease in dynamic tubular events (Fig. 4F-G).

**Figure 4.**
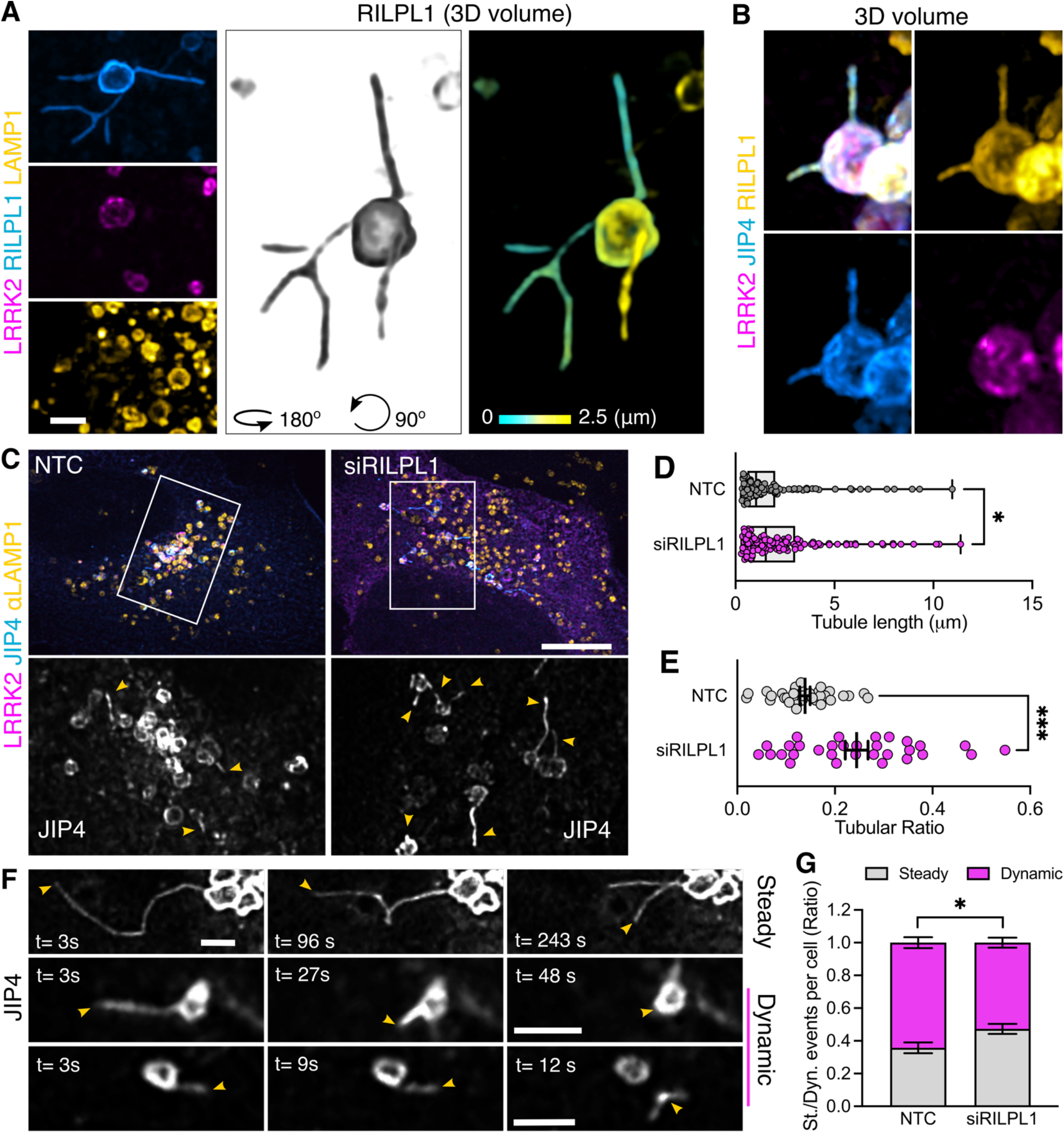
**RILPL1 colocalizes with LYTL tubules and reduces tubulation**. (A) Mouse primary astrocytes were transfected with HaloTag-LRRK2, mCherry-RILPL1 and mNeonGreen-LAMP1. 48 h later, cells were treated with LLOME for 6 h and imaged live. The 3D volume shows different RILPL1 tubules emanating from a lysosome. (B) U2OS cells were transfected with HaloTag-LRRK2, mCherry- RILPL1 and mNeonGreen-JIP4. 36 h later, cells were treated with LLOME for 2 h and fixed. 3D volume view of a lysosome with two LYTL tubules expressing JIP4 and mCherry. (C) U2OS cells were treated with NTC or siRILPL1 for 24 h. Then cells were transfected with HaloTag-LRRK2 and mNeonGreen-JIP4 for 36 h. Cells were treated with LLOME for 2 h, fixed and stained with an anti-LAMP1 antibody. (E) Box plot showing the tubule length in both groups. Unpaired t-test was applied (p=0.0458, n= 119-134 total tubules analyzed respectively). Data are mean ± SEM. (F) Tubular ratio quantification. Data are mean ± SEM (p=0.0002, n = 31-33 cells). Unpaired t-test with Welch’s correction was used. (G) U2OS cells were treated with NTC or siRILPL1 for 24 h. Then cells were transfected with 3xflag- LRRK2 and mNeonGreen-JIP4 for 36 h. Cells were treated with LLOME for 2 h and analyzed live. Picture depicts examples of “steady” and “dynamic” tubular events. Graph showing the ratio of steady and dynamic tubular events per cell in the different conditions. Data are mean ± SEM (p=0.0141, n = 20 cells). Unpaired t-test. Scale bar (D)= 10 µm; (A,C,G)= 2 µm.

### RILPL1 binds to p150^Glued^ and drives organelle movement to the minus-end of microtubules

A common feature of the RHD family members is their role as motor adaptor proteins. JIP3 and JIP4 can bind to dynein/dynactin and kinesins, while RILPL2 binds to myosin-Va. Although RILPL1 has not previously been associated with any motor, *in vitro* work has proposed a potential interaction between RILPL1 and dynein-light intermediate chain (DLIC) (Celestino et al., 2022). As dynein/dynactin move organelles towards the centrosome, we asked whether RILPL1 mediates its function on LRRK2-positive lysosomes by acting as a dynein adaptor protein. This hypothesis would be consistent with the observation that LYTL tubules elongated towards the plus-end of the microtubules, away from the centrosome (Fig. 5A-C), further suggesting that LYTL tubules elongate via kinesin and retract through dynein/dynactin. Given that JIP3 and JIP4 bind to both DLIC and to the dynactin subunit p150^Glued^ (DCTN1) (Vilela et al., 2019), we assessed if RILPL1 is able to bind to these proteins. Immunoprecipitation assays reveal that RILPL1 binds to p150^Glued^, although we could not reproduce the RILPL1: DLIC interaction in our experimental conditions (Fig. 5D).

**Figure 5.**
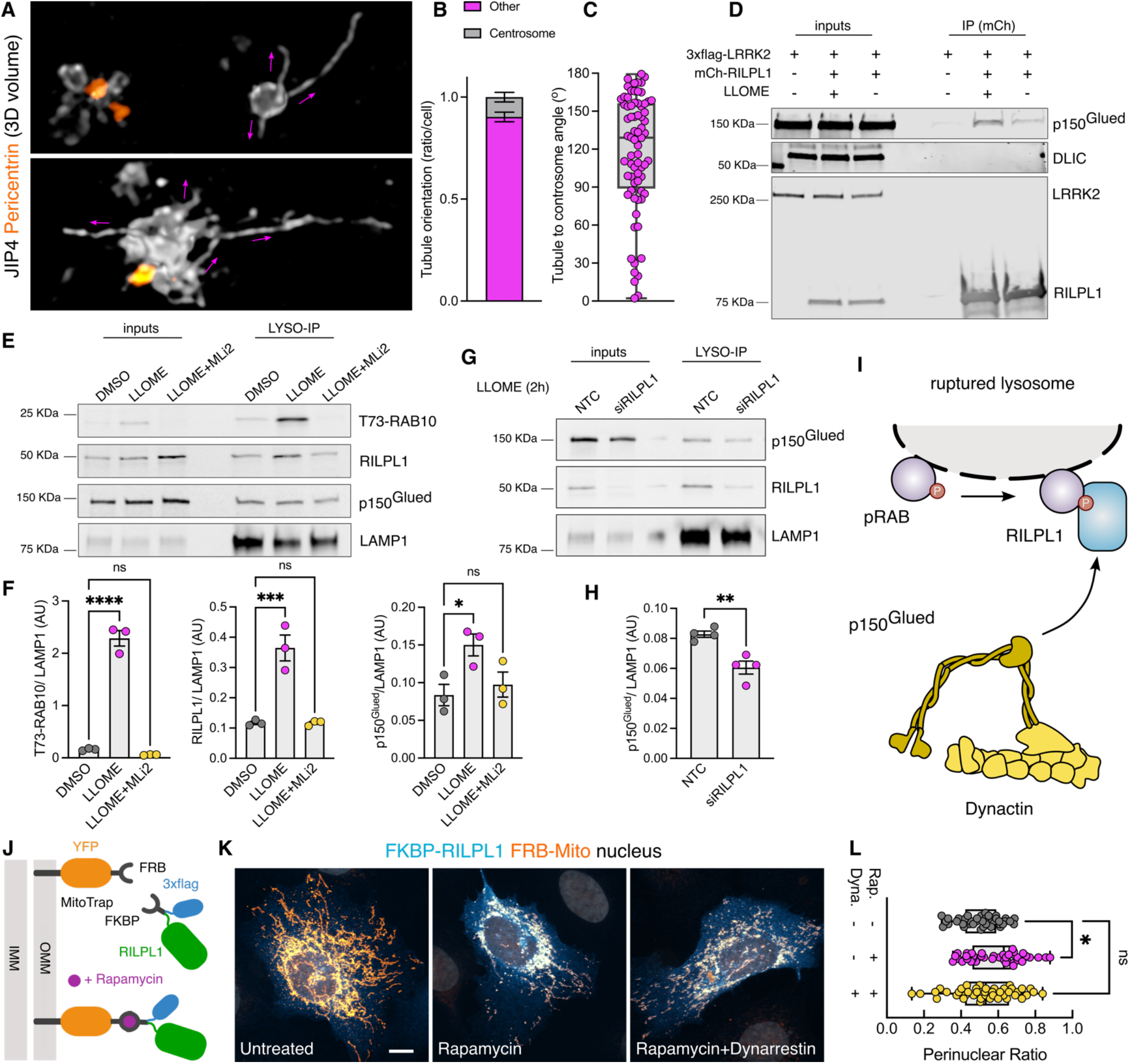
RILPL1 binds to p150^Glued^ and favors its recruitment to lysosomes. (A) U2OS cells were transfected with HaloTag-LRRK2 and mNeonGreen-JIP4 for 36 h. Cells were then treated with LLOME (2 h), fixed and stained with pericentrin. Volume view image of JIP4+ tubules and centrosomes (pericentrin). (B) The orientation of the JIP4+ tubules were manually annotated as “towards the centrosome” (centrosome) or “elsewhere” (other) (n=28 tubules). (C) Box plot depicting the tubule-to-centrosome angle (n= 72 tubules from 14 cells). (D) HEK293FT cells were transfected with 3xflag-LRRK2 and mCherry-RILPL1 for 24 h. Cells were treated or not with LLOME for 2 h and lysates were subjected to immunoprecipitation with anti-RFP antibodies. (E) HEK293T cells stably expressing GFP- LRRK2 and TMEM192-3xHA were seeded and treated with DMSO, LLOME or LLOME + MLi2 24 h later. Lysosomes were purified with anti-HA beads following the LYSO-IP technique. Lysosomes were then lysed, and their content was analyzed via immunoblotting. (F) Quantification of T73-RAB10, RILPL1 and p150^Glued^ protein levels from the lysosomal fraction. Data are mean ± SEM from three independent replicates. p(T73-RAB10)<0.0001; p(RILPL1)= 0.0008; p(p150^Glued^)= 0.0368. One-way ANOVA with Dunnett’s. (G) HEK293T cells stably expressing GFP-LRRK2 and TMEM192-3xHA were seeded and treated with NTC and siRILPL1 for 48 h later. Then, cells were treated with LLOME for 2 h. Lysosomes were purified with anti-HA beads following the LYSO-IP technique. Lysosomes were then lysed, and their content was analyzed via immunoblotting. (H) Quantification of p150^Glued^ protein levels from the lysosomal fraction. Data are mean ± SEM from four independent replicates. p= 0.0036. Unpaired t-test. (I) Schematic representation of our experimental model where the pRAB proteins recruit RILPL1, which in turn binds to p150^Glued^ and recruits it to membrane damaged lysosomes. (J) Cartoon explaining the FRB-FKBP system to trap RILPL1 to the outer mitochondrial membrane (OMM). (K) U2OS cells were transfected with Mito-YFP-FRB and 3xflag-FKBP-RILPL1 for 24 h. Cells were treated or not with rapamycin, or rapamycin+dynarrestin. (L) Box plot showing the perinuclear ratio in all groups. Data are mean ± SEM (p= 0.0135, n = 39-42 cells). One- way ANOVA with Dunnett’s post hoc test. Scale bar= 10 µm.

Our mass spectrometry data on isolated lysosomes shows a LRRK2 kinase-dependent recruitment of three cytoplasmic dynein subunits, the dynein activator LIS1, and four dynactin complex members, including p150^Glued^ (Fig. S1C). We validated this result using western blotting (Fig. 5E-F), demonstrating that LRRK2 kinase activity is required to recruit p150^Glued^ to damaged lysosomes in a similar pattern than RILPL1. We then reasoned that LRRK2 kinase activity recruits p150^Glued^ through RILPL1. RILPL1 knockdown significantly reduces the amount of p150^Glued^ on lysosomes in LLOME-treated cells (Fig. 5G-H), suggesting that RILPL1 is responsible for the recruitment of dynactin to damaged lysosomes (Fig. 5I). To address whether RILPL1 is sufficient to promote transport of organelles towards the nucleus, we used the FKBP-FRB system to localize RILPL1 to mitochondria by co-expressing a Mito-YFP-FRB construct and a 3xflag-FKBP-RILPL1 vector (Fig. 5J). In the presence of rapamycin, FRB and FKBP dimerize and RILPL1 translocates to the mitochondria outer membrane (Fig. 5J-K). This leads to a significant clustering of the mitochondria towards the perinuclear area, reversed by pre-treatment with the dynein inhibitor Dynarrestin (Fig. 5K-L). Our data suggest that RILPL1 can act as a dynein adaptor protein through its interaction with p150^Glued^, leading to clustering of LRRK2-positive lysosomes towards the centrosome and retraction of LYTL tubules.

### LYTL tubules elongate through tyrosinated microtubules

Motor proteins move organelles through microtubule tracks. We have previously shown that nocodazole disrupts JIP4-positive tubule formation, demonstrating that microtubules are important in tubule elongation. We confirmed the association between LYTL tubules and microtubules in live cells (Fig. S4A; Suppl. Movie 2), showing a tubule elongating on a microtubule (Fig. S4A, outline arrowhead; open arrowhead on Suppl. Movie 2). ɑ-Tubulin undergoes critical post-translational modifications (PTMs) that can regulate microtubule functions (for reviews, see (Nieuwenhuis and Brummelkamp, 2019; Roll-Mecak, 2020; Janke and Magiera, 2020). Among them, acetylation on lysine 40 residue and tyrosination/detyrosination on its C-terminal tail is particularly important. Indeed, ɑ-tubulin PTMs regulate the efficiency of motors, as motors have preference for certain PTMs. For example, dynein has preference for tyrosinated microtubules and kinesin-1 family members prefer acetylated/detyrosinated microtubules. Acetylated ɑ-tubulin is largely detyrosinated (Dunn et al., 2008; Katrukha et al., 2021) (Fig. S4B), which allows the separation of most microtubules into two main groups: acetylated and tyrosinated (Tas et al., 2017; Guardia et al., 2016; Katrukha et al., 2021) (Fig. S4B).

We therefore considered whether tubulin PTMs could affect tubule elongation. When analyzed in fixed cells, LYTL tubules show greater association with tyrosinated microtubules than acetylated microtubules (Fig. 6A-C; Fig. S4C), and live-cell imaging experiments show a JIP4-positive tubule growing on a tyrosinated microtubule (Fig. 6D; Fig. S4D). The sorted material also travels through microtubules (Fig. S4E; Suppl. Movie 3), and its movement is hampered by nocodazole (Fig. S4F; Suppl. Movie 4). Time-lapse microscopy shows a sorted tubule traveling on a tyrosinated microtubule (Fig. E; Suppl. Movie 5). ɑ-tubulin tyrosination depends on tubulin tyrosine ligase (TTL) (Fig. 6F) (Szyk et al., 2011) and it is known that TTL depletion leads to a dramatic increase in detyrosinated ɑ-tubulin (Peris et al., 2006; Zink et al., 2012; Peris et al., 2009; Nieuwenhuis et al., 2017) (Fig. S5A-D). TTL depleted cells displayed lesser lysosomal tubulation than control cells (Fig. 6G-H), consistent with the effect of nocodazole blocking tubulation (Bonet-Ponce et al., 2020). Nocodazole addition depolymerizes the microtubule network leaving few nocodazole-resistant microtubules (Xu et al., 2017) (Fig. S5E) which are acetylated/detyrosinated and not tyrosinated (Xu et al., 2017; Kesarwani et al., 2020) (Fig. S5F).

**Figure 6.**
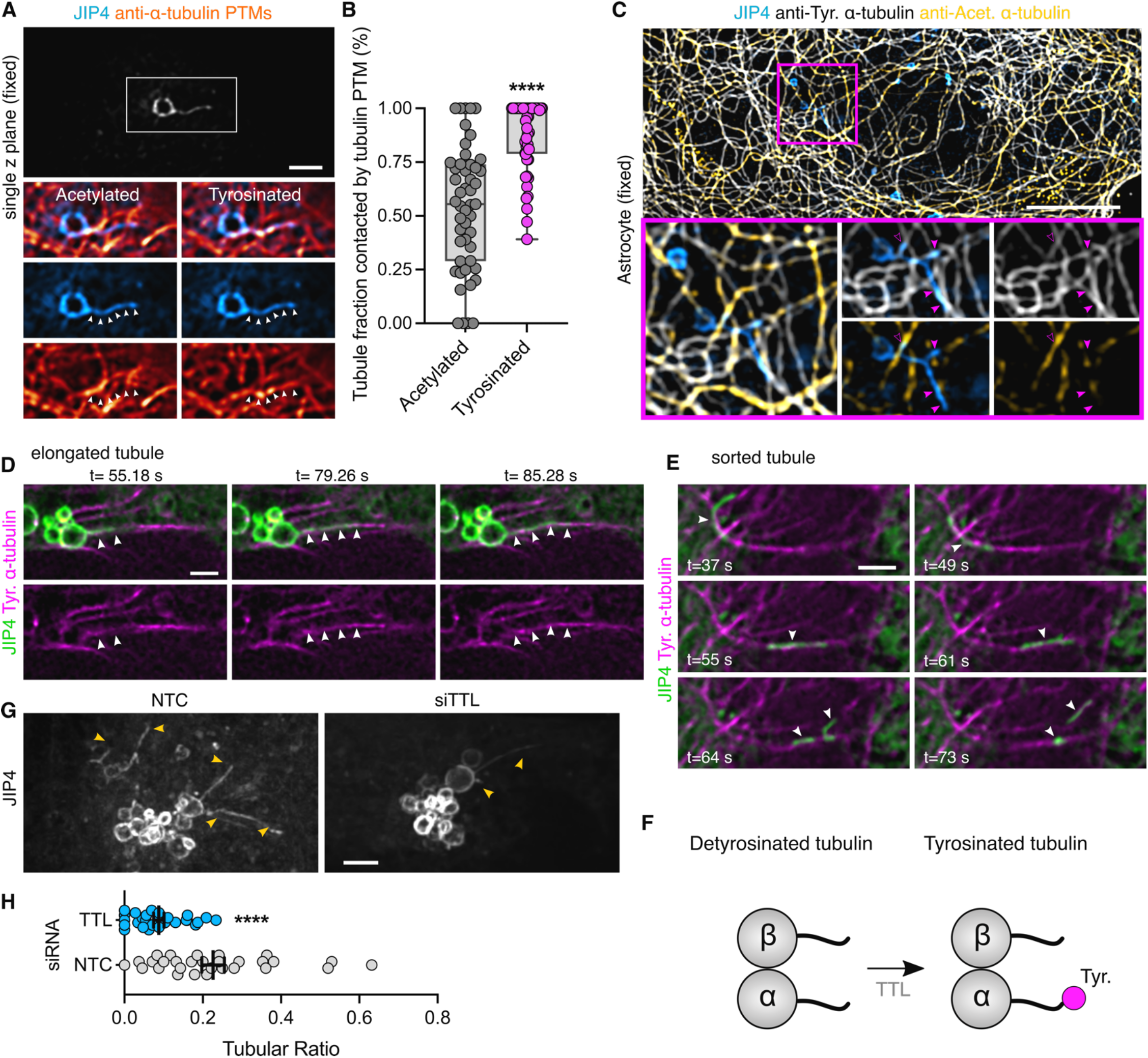
LYTL tubules elongate through tyrosinated microtubules. (A) U2OS cells were transfected with 3xflag-LRRK2 and GFP- JIP4. Cells were treated with LLOME (2 h) and fixed. After fixation cells were stained for GFP, tyrosinated ɑ-tubulin and acetylated ɑ- tubulin. (B) Box plot showing the LYTL tubule contact fraction with the different ɑ-tubulin PTMs. Unpaired t-test with Welch’s correction was applied (p<0.0001, n= 49 tubules). Arrowheads show lysosomal tubules. Scale bar= 2 µm. (C) Mouse primary astrocytes were transfected with 3xflag-LRRK2 and GFP-JIP4. Cells were treated with LLOME (6 h) and fixed. After fixation cells were stained for GFP, tyrosinated ɑ-tubulin and acetylated ɑ-tubulin. (D-E) Time-lapse experiment of U2OS cells transfected with 3xflag-LRRK2, mNeonGreen- JIP4 and TagRFP-T-A1aY1 (tyrosinated microtubules) and treated with LLOME for 2 h before imaging. (F) The enzyme TTL is responsible for adding the tyrosine residue into the C terminal tail of ɑ-tubulin. (G) U2OS cells were incubated with a non-targeting control (NTC) or TTL siRNA and transfected 24 h later with 3xflag-LRRK2 and mNeonGreen-JIP4. Cells were treated with LLOME (2 h) and imaged live. The tubular ratio was quantified. (D) Graph depicts the tubular ratio in the NTC and siTTL groups. Unpaired t-test with Welch’s correction was applied. Data are mean ± SEM (p<0.0001, n= 27-29 cells). Magenta arrowheads show colocalization between LYTL tubules and only tyrosinated microtubules (filled) and tyrosinated + acetylated microtubules (empty). White arrowheads in (A) show the length of a LYTL tubule; in (D), show a budding tubule elongating on a tyrosinated microtubule; and in (E), a sorted tubule traveling to a tyrosinated microtubule. Yellow arrowheads indicate JIP4-positive tubules. Scale bar (A,C)= 10 µm; (D-G)= 2 µm.

Taken together, our data show that LYTL tubules move through tyrosinated microtubules, which is consistent with the observation that the dynein/dynactin complex requires ɑ-tubulin tyrosination to initiate their retrograde movement (McKenney et al., 2016; Nirschl et al., 2016). As LYTL tubule retraction is mediated by the RILPL1: dynein/dynactin complex, tubule elongation to the plus-end of microtubules is likely driven by JIP4 and kinesin(s). Even though JIP4 has long been associated with KIF5B (a kinesin-1 family member), KIF5B knockdown has no effect on tubulation (Fig. S5G), suggesting that another kinesin is responsible for tubule elongation. Our data is consistent with the fact that kinesin-1 family members preferentially move on acetylated/detyrosinated microtubules. It is therefore likely that the kinesin(s) associated with JIP4 in LYTL tubule elongation have preference for tyrosinated microtubules (Fig. 7).

**Figure 7.**
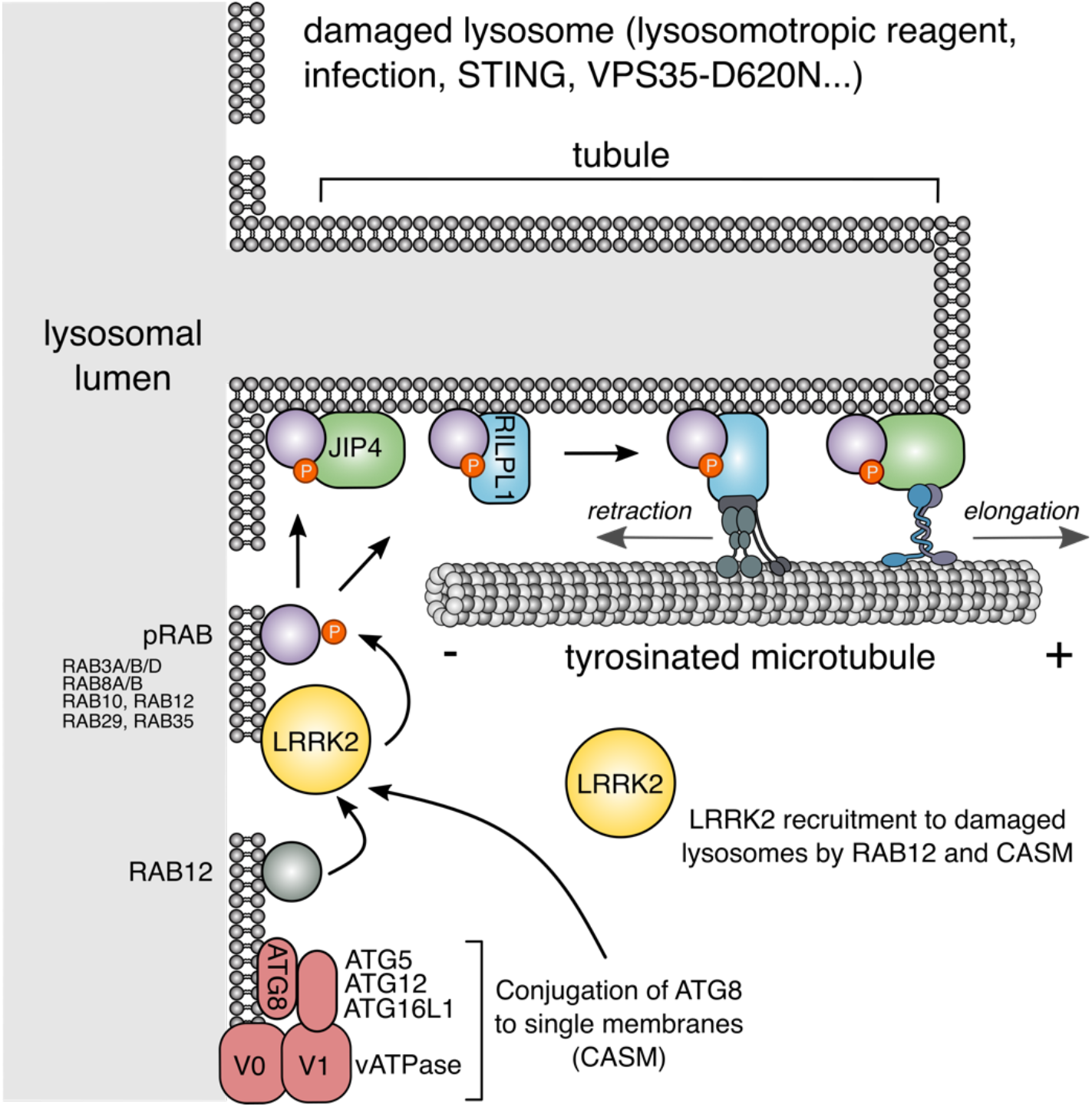
Schematic representation of our working model. LRRK2 is recruited to ruptured lysosomes (likely via RAB12 and CASM), where it phosphorylates and recruits RAB substrates. pRABs recruit their effectors JIP4 and RILPL1. JIP4, through an as yet unknown kinesin, elongates the LYTL tubules, whereas RILPL1 via dynein/dynactin, favors tubule retraction.

## DISCUSSION

Lysosomes change the shape of their membrane to adapt to various cellular responses and undergo distinct mechanisms of membrane tubulation. For example, in the context of immune cell response to LPS treatment, lysosomes undergo a transformation from vesicular to tubular morphology, a change essential for MHC-II presentation (Chow et al., 2002; Saric et al., 2016). When the lysosomal pool becomes saturated, lysosomes generate new lysosomes through a process known as Lysosomal Reformation (LR) (Yu et al., 2010). In LR, new lysosomes emerge through the extrusion of membranes from pre-existing healthy lysosomes, featuring membrane markers such as LAMP1, and this process is facilitated by the motor protein KIF5B (Du et al., 2016). We have previously described LYTL as a novel lysosomal tubulation and vesicle sorting process that is orchestrated by the Parkinson’s disease kinase LRRK2. LYTL originates from compromised lysosomes and likely serves as a mechanism for transferring undegraded cargo from dysfunctional lysosomes to active ones.

In this study, we systematically mapped the LRRK2 lysosomal proteome. Notably, our screening highlighted RAB proteins as the most abundant group, confirming prior data that these proteins are the primary targets of active LRRK2 on lysosomes and underscores the reliability of our screening approach. Among the proteins identified, two potential pRAB effectors, JIP4 and RILPL1, were observed. Given their roles as motor adaptor proteins in LYTL, it is reasonable to consider that LYTL is the major consequence of accumulation of LRRK2 on dysfunctional lysosomes. It is also worth noting that calcium-binding proteins and RNA-binding proteins were also hits in this screen and may be of interest to follow up in additional studies.

RILPL1 has been previously linked to LRRK2 in other cellular compartments including the centrosome and the primary cilia (Lara Ordónez et al., 2019; Dhekne et al., 2018). It has been proposed that hyperactive LRRK2 disrupts centrosomal cohesion and prevents ciliogenesis via pRABs and RILPL1. In agreement with our results, recent work also demonstrates a LRRK2 dependent recruitment of RILPL1 to lysosomes, in cells harboring the VPS35-D620N mutation (Pal et al., 2023). However, the molecular function of RILPL1 has not been clarified. Here we show that RILPL1 clusters LRRK2-positive lysosomes around the centrosome and reduces LYTL- dependent tubulation. RILPL1 binds to p150^Glued^, the largest subunit of dynactin which binds to both microtubules (Waterman-Storer et al., 1995) and cytoplasmic dynein (Vaughan and Vallee, 1995), promoting retrograde transport of cargo to the minus-end of microtubules (Waterman-Storer et al., 1997). Our data demonstrate that RILPL1 promotes lysosomal retrograde transport and tubule retraction via the dynein/dynactin complex. The possibility that RILPL1 might act as a motor adaptor protein aligns with the observed role of the other members of the RHD family, all of which are known to share this function. Our data suggests a very tight regulation of LYTL dynamics, with both motor adaptor proteins working in opposite directions.

The dynein/dynactin complex relies on tubulin tyrosination to initiate retrograde movement (McKenney et al., 2016; Nirschl et al., 2016) and, as shown here for the first time, LYTL tubules extend along tyrosinated microtubules. Since JIP4 is essential for promoting the elongation of LYTL tubules towards the plus-end of microtubules, it is plausible that JIP4 interacts with kinesin proteins to facilitate this process. When JIP4 is recruited to membranous compartments by Arf6, JIP4 preferentially binds to dynein/dynactin (Montagnac et al., 2009; Cason and Holzbaur, 2023). However, our data suggests that once JIP4 is recruited to the lysosomal surface by pRABs, JIP4 binds to kinesin. This observation aligns with the other studies of the role of pRABs and JIP4 in neuronal autophagosomal transport (Boecker et al., 2021; Dou et al., 2023; Cason and Holzbaur, 2023). Although JIP4 has primarily been associated with members of the kinesin-1 family, knockdown of KIF5B did not result in a decrease in LYTL tubulation. This finding suggests that KIF5B does not play a role in tubule elongation, and that JIP4 binds to a different kinesin in this model, consistent with the fact that kinesin-1 family members travel on acetylated/detyrosinated microtubule tracks. The kinesin(s) responsible for tubule elongation are therefore still unknown and will be addressed in future work.

In summary, here we have explored the role of RILPL1 once recruited by LRRK2 to dysfunctional lysosomes. We describe new regulatory proteins of LYTL, which is likely pathologically relevant given the role of LRRK2 and lysosomes in PD.

## ACKNOWLEDGEMENTS

This research was supported by the Intramural Research Program of the NIH, National Institute on Aging (MRC). All the data needed to evaluate the conclusions in the paper are present in the paper and/or the Supplementary Materials. Additional data related to this paper may be requested from the authors.

## AUTHOR CONTRIBUTIONS

Conceptualization, L.B-P., M.R.C.; Methodology, L.B-P., Y.L., J.H.K., A.B., T.T.; Formal Analysis, L.B-P., M.R.C.; Investigation, L.B-P., Y.L., J.H.K., A.B., T.T; Writing, L. B-P.; Supervision, M.R.C., L.B-P.; Funding, M.R.C.

## METHODS

### Cell culture

U2OS cells (ATTC), HEK293FT, HEK293T, and mouse primary astrocytes were maintained in DMEM (Thermo Fisher) containing 4.5 g/l glucose, 2 mM l-glutamine, and 10% FBS (Gibco) at 37°C in 5% CO2 into 75 cm^2^ tissue culture flasks. All cell lines were used until passage 30. Cells were seeded on 12 mm coverslips for fixed experiments and in 35 mm glass bottom dishes (MatTek) for live-cell imaging experiments. Coverslips and dishes were pre-coated with matrigel (Corning).

Primary astrocyte cultures were prepared from C57BL/6J newborn (postnatal day 0) pups. Dissected mouse cortices were incubated in 1 ml/cortex Basal Medium Eagle (BME) (Sigma-Aldrich), containing 5 U of papain/(Worthington) for 30 min at 37°C. Five micrograms of deoxyribonuclease I was added to each cortex preparation, and brain tissue was dissociated into a cellular suspension that was washed twice with 10 volumes of BME and counted. Astrocyte cultures were plated in Dulbecco’s modified Eagle’s medium (DMEM) (Thermo Fisher Scientific), supplemented with 10% fetal bovine serum (FBS) (Gibco) into 75 cm^2^ tissue culture flasks. For the preparation of purified astrocyte cultures, 7- to 10-day primary cultures were vigorously shaken to detach microglia and oligodendrocytes.

The HEK293T-inducible GFP-LRRK2-WT cell line was obtained from D. Alessi (University of Dundee), and expression was induced by addition of doxycycline (Nichols et al., 2010).

### Reagents and treatments

LLOME (Sigma-Aldrich, L7393) was diluted in dimethyl sulfoxide (DMSO) and added at 1 mM for the indicated times. MLi-2 (Tocris, 5756; and MedChemExpress, HY-100411) was used at 1 μM in DMSO for 90 min prior to LLOME incubation. Nocodazole (Sigma-Aldrich, M1404) was diluted in DMSO and added at 10 μM, 30 min before imaging. HaloTag–transfected cells were incubated with the JFX650 peptide (Janelia) at 100 nM for 10 or 40 min, and cells were then washed three times and incubated with fresh medium before treated with LLOME. For experiments utilizing the FKBP-FRB complex, rapamycin (Cayman Chemicals, 3346) was added at 200 nM for 2 h prior to fixation in 4% PFA. Dynarrestin (Tocris, 6526) was added at 25 μM, 30 min before rapamycin.

### Antibodies

The following primary antibodies were used: mouse anti-FLAG M2 (Sigma-Aldrich, F3165; 1:500 for immunocytochemistry (ICC) and 1:10,000 for western-blot (WB)), rabbit anti-JIP4 (Cell Signaling Technology; 5519, 1:1000 for WB and 1:100 for ICC), rabbit anti-LAMP1 (Cell Signaling Technologies, D2D11; 1:2000 for WB and 1:200 for ICC), rat anti-LAMP1 (Developmental Studies Hybridoma Bank (DSHB), 1D4B; 1:100 for ICC), mouse anti-LAMP1 (DSHB, H4A3; 1:1000 for WB and 1:100 for ICC), mouse anti-LAMP2 (DSHB, H4B4; 1:1000 for WB and 1:100 for ICC), mouse anti-p150^Glued^ (BD Biosciences, 610474; 1:10,000 for WB), rabbit anti-DLIC (Bethyl Laboratories, A304-208A; 1:1000 for WB), chicken anti-GFP (Aves Lab, GFP-1020; 1:500 to 1:1000 for ICC), mouse anti-GFP (Roche, 11814460001; 1:10,000 for WB), mouse anti-α-tubulin (Cell Signaling Technology, 3873; 1:10,000 for WB), rat anti-tyrosinated ⍺-tubulin (YL1/2) (Abcam, ab6160; 1:10,000 for WB and 1:500 for ICC), rabbit anti-detyrosinated ⍺-tubulin, Abcam, ab48389; 1:1000 for WB and 1:100 for ICC), mouse anti-acetylated ⍺-tubulin (Millipore-Sigma, T7451: 1:200 for ICC), rabbit anti-total ⍺-tubulin (Abcam, ab52866; 1:10,000 for WB), rabbit anti-RILPL1 (Abcam, MJF-R41-21 (ab302492); 1:100 for ICC and 1:1000 for WB), rabbit anti-RAB10 (Abcam, ab237703; 1:100 for ICC and 1:2000 for WB), rabbit anti-RAB10 (phospho- T73) (Abcam, MJF-R21-22-5 (ab241060); 1:100 for ICC and 1:2000 for WB), rabbit anti-LRRK2 (Abcam, ab133474; 1:1000 for WB), rat anti-HA (7C9) (Chromotek, 7c9-100; 1:10,000 for WB and 1:500 for ICC), rabbit anti-EEA1 (Cell Signaling Technology, 3288; 1:100 for ICC), rabbit anti-LC3B (Cell Signaling Technology, 2775; 1:1000 for WB), rabbit anti-LAMTOR4 (Cell Signaling Technologies, 12284; 1:1000 for WB), rabbit anti-calreticulin (D3E6) (Cell Signaling Technologies, 12238; 1:1000 for WB), mouse anti-GM130 (H7) (Santa Cruz Biotechnology, sc-55590; 1:1000 for WB), rabbit anti-GAPDH (Millipore-Sigma, G9545; 1:10,000 for WB), rabbit anti-Cathepsin D (Millipore, 219361, 1:2000 for WB), mouse anti-pericentrin (Abcam, ab28144; 1:100 for ICC) and rabbit anti-Tom20 (FL-145) (Santa Cruz Biotechnology, sc-11415; 1:5000 for WB).

For ICC, unless otherwise stated, the secondary antibodies were purchased from Thermo Fisher Scientific. The following secondary antibodies were used: donkey anti-mouse Alexa Fluor 568 (A10037, 1:500), donkey anti-rabbit Alexa Fluor 488 (A-21206, 1:500), donkey anti-mouse Alexa Fluor 568 (A-21202, 1:500), donkey anti-rat Alexa Fluor 488 (A-21208, 1:500), donkey anti-rabbit Alexa Fluor 568 (A10042, 1:500), donkey anti-mouse Alexa Fluor 647 (A-31571, 1:500), and goat anti-rat Alexa Fluor 647 (A-21247, 1:250 to 1:500). Donkey anti-chicken Alexa Fluor 488 (703-545-155, 1:500) and donkey anti-rat Alexa Fluor 405 (712-475-153, 1:100) were obtained from Jackson ImmunoResearch.

### Plasmids

Constructs for 3xFLAG-LRRK2, HaloTag-LRRK2, mScarlet-LRRK2, LYSO-LRRK2, mNeonGreen-JIP4 and HaloTag-JIP4 have been described previously (Beilina et al., 2014, 2020; Bonet-Ponce et al., 2020; Bonet-Ponce and Cookson, 2022b; Kluss et al., 2022a). RILPL1 cDNA was amplified with PCR and cloned into the pCR8/GW/TOPO vector (Thermo Fisher). PCR8-RILPL1 was then subcloned into pDEST-3xflag-FKBP using Gateway technology (Thermo Fisher). mCherry-RILPL1 was created with IN-FUSION (Takara) from the mCherry-Climp63 vector (Addgene#136293) (Shibata et al., 2008). RILPL1-R293A was created using the QuickChange Lightning Site-directed mutagenesis kit (Agilent) from the mCherry-RILPL1 (WT) vector.

LAMP1-RFP, LAMP1-mNeonGreen, TMEM192-3xHA, LifeAct-mNeonGreen, 6xHis-TagRFP-T_A1aY1, pEYFP- Mitotrap, TUBB5-Halo, EMTB-mNeonGreen, plasmids were purchased from Addgene (Addgene#1817, Addgene#98882, Addgene#102930, Addgene#98877, Addgene#158754, Addgene#46942, Addgene#6469, Addgene#137802) (Sherer et al., 2003; Chertkova et al., 2017; Abu-Remaileh et al., 2017; Kesarwani et al., 2020; Robinson et al., 2010; Uno et al., 2014).

### Transfection

Transient transfections of U2OS and mouse primary astrocytes cells were performed using Lipofectamine Stem Reagent and Opti-MEM (Thermo Fisher). HEK293FT were transfected using Lipofectamine 2000 and Opti-MEM (Thermo Fisher). HEK293FT cells were transfected for 24 h, U2OS cells were transfected for 36-48 h prior to imaging. Astrocytes were transfected for 48 h prior to imaging. For siRNAs, cells were transfected with the SMARTpool ON-TARGET (Horizon) plus scramble, or TTL, RILPL1 and KIF5B siRNA using Lipofectamine RNAiMAX (Thermo Fisher) transfection reagent. U2OS cells were incubated with siRNA for a total of 3 days before imaging or lysis. HEK293T cells were incubated with siRNA for 48 h prior to performing the LYSO-IP.

### GFP-LRRK2/ TMEM193-3xHA Stable cell production

HEK293T cells stably expressing GFP-LRRK2 were transfected with TMEM192-3xHA, using Lipofectamine 2000 at a 75% confluency in a 75 cm^2^ flask. 24 h later, cells were treated with puromycin (1 μg/ml) for 7 days. The polyclonal population was seeded in a density of one cell per well in 96 well plates. The presence of TMEM192-3xHA was corroborated by staining and immunoblotting in the selected clone.

### Lysosome Immunoprecipitation (LYSO-IP)

Lysosomes from HEK293T cells stably expressing GFP-LRRK2 and TMEM192-3xHA were purified following the protocol described in (Davis et al., 2021). Briefly, cells were seeded in 15 cm dishes (proteomics) or 10 cm dishes (immunoblotting) at a density appropriate for them to reach confluency after 24h or 48h (for siRNA experiments). All subsequent steps were performed on ice or at 4°C unless otherwise noted. After media removal, cell monolayers were rinsed with ice-cold KPBS buffer (136 mM KCl, 10m M KH2PO4, pH 7.25, supplemented with 1x protease and phosphatase inhibitor cocktail (Thermo Fisher)), scraped into 10 ml (proteomics) or 1 ml (immunoblotting) of KPBS and collected by centrifugation at 300 g for 5 min. Pelleted cells were resuspended in a total volume of 1 ml KPBS (supplemented with 3.6% (w/v) OptiPrep (Sigma)) and fractionated by passing through a 23 G syringe five times followed by centrifugation at 700 g for 10 min. Post-nuclear supernatant was harvested and incubated with 100 μl (proteomics) or 40 μl (immunoblotting) of anti-HA magnetic beads (Thermo Fisher, prewashed with KPBS buffer) with end-over-end rotation for 15 min at 4°C. Lysosome-bound beads were washed two times with KPBS(+ OptiPrep) and two times with KPBS. Samples were incubated with lysis buffer (20 mM tris-HCl (pH 7.5), 150 mM NaCl, 1 mM EDTA, 0.3% Triton X-100, 10% glycerol, 1× Halt protease and phosphatase inhibitor cocktail) for 25 min with end-over-end rotation at 4°C. Beads were removed and lysates were snap-frozen with LN2 (proteomics), or boiled at 95°C for 5 min with 4x loading buffer (Biorad) and β-mercaptoethanol (immunoblotting).

### Mass spectrometry Analysis

In-gel samples were reduced with 5 mM tris(2-carboxyethyl)phosphine (TCEP) (Sigma Aldrich), alkylated with 5 mM N-Ethylmaleimide (NEM), (Sigma Aldrich). Samples were digested with trypsin (Promega) at 1:10 (w/w) trypsin:sample ratio at 37C for 18 hr. Peptides were extracted then desalted using Oasis HLB plate (Waters). An Orbitrap Lumos mass spectrometer (Thermo Scientific) coupled in line with an Ultimate 3000 HPLC (Thermo Scientific) was used for data acquisition. Peptides were directly injected and separated on a ES803A column (75-μm inner diameter, 25 cm length, 2 μm C18 beads; Thermo Scientific) using a gradient with mobile phase B (0.1% formic acid in LC-MS grade acetonitrile) increased from 5 to 25% in 90 min at a flow rate of 300 nl/min. LC-MS/MS data were acquired in data dependent mode. The survey scan was performed at a resolution of 120k, and a mass range of 400–1500 m/z. MS2 data were acquired in ion trap with rapid scan rate and an isolation window of 1.6 m/z. CID method with a fixed collision energy of 30 was used for peptide fragmentation. The minimum signal intensity required to trigger MS2 scan was 1e4.

Raw data were process using Proteome Discoverer 2.4 Software. Data were searched against the Uniprot Human database. The mass tolerances for precursor and fragment were set to 5 ppm and 0.6 Da, respectively. Up to 1 missed cleavage was allowed. NEM on cysteines was set as fixed modification. Variable modifications include Oxidation (M), Met-loss (Protein N-term) and Acetyl (Protein N-term). 1% false discovery rate (FDR) at protein level was applied. Proteins detected with 1 peptide were further filtered out from the results. The protein abundances were obtained by summing abundance of the connected peptides, then normalized to the total peptide amount. Protein ratios were calculated by comparing the median protein abundance of each group. ANOVA (Individual Proteins) method was used for hypothesis test.

### Confocal microscopy

#### Airyscan

Airyscan images were taken using a Zeiss LSM 880 microscope equipped with a 63X 1.4 NA objective. Super-resolution imaging was performed using the Airyscan mode. Raw data were processed using Airycan processing in ‘auto strength’ mode with Zen Black software version 2.3.

#### SoRa

For spinning disk super-resolution microscopy we used a W1-SoRa super-resolution spinning disk microscope (Nikon) with a 60X 1.49 NA oil immersion objective. A 2.8X intermediate magnification (168X combined) was used in time-lapse experiments regarding JIP4 tubules. A 4X intermediate magnification (240X combined) was used in snapshot images. An offset micro-lensed SoRa disk was used and an environmental chamber to maintain cells at 37°C with humidified 5% CO2 gas during imaging. For deconvolution, we used 20-25 iterations of the 3D Landweber algorithm with the NIS-Elements AR 5.21.03 software. Images from two channels were acquired simultaneously using a Cairn twin-cam emission splitter and two Photometrics prime 95b sCMOS cameras, a 565LP DM, and appropriate emission cleanup filters. Triggered piezo was used to maximize speed. Stacks were taken with 0.2 µm distance between slices.

When needed, bleaching was corrected using the “Histogram matching” option from Fiji (ImageJ). Unless otherwise stated, stacks were processed as maximum intensity projection.

### Lookup tables (LUTs)

All pseudocolors used in this paper can be found in the “NeuroCyto LUTs” collection (Fiji, ImageJ). For gray colors, “JDM grays” were used. For cyan colors, “cyan”, “cyan hot” and “JDM04 pop cyan” were used. For magenta, “Magenta” and “Magenta Hot” were used. For orange, “NanoJ-Orange” was used. For yellow, “Yellow Hot” was used. For inverted LUTs, “JMD, gamma inverted” were used. For z color code, “Z-stack Depth Colorcode 0.0.2” plugin (Fiji, ImageJ) was applied using “DavLUT-Bright” as LUT.

### Volume view

Volume view was obtained using the NIS-Elements AR 5.21.03 software (Nikon) with the “Alpha Blending” option with or without “Depth Color Code”. Volume view was also obtained with the “volume viewer” plugin from Fiji (ImageJ, NIH).

### Tubular ratio measurements

U2OS cells were transfected with the 3xflag-LRRK2 plasmid, and co-transfected with mNeonGreen-JIP4. After treatment with LLOME, live cells were imaged with SoRa. The tubular ratio in each cell was measured as: Ratio= #JIP4+tubules/ #JIP4+lysosomes. Only cells with 10 or more JIP4-positive lysosomes per cell were imaged.

### LYTL tubule contact to ɑ-tubulin PTM

U2OS cells were transfected with 3xflag-LRRK2 and pDEST53-JIP4 for 36 h. After treatment with LLOME for 2 h, cells were fixed and stained for GFP, tyrosinated ɑ-tubulin and acetylated ɑ-tubulin. “Tubule fraction contacted by ɑ-tubulin PTM” was measured as: distance of contact between LYTL tubule and ɑ-tubulin PTM/ total distance of LYTL tubule.

### Perinuclear Ratio

LAMP1 and Mito-YFP integrated densities were measured using Fiji (ImageJ, NIH) after thresholding. Every cell was thresholded according to its individual intensity, capturing the whole lysosomal or mitochondrial population. Integrated density was measured for the whole cell (Total), and the area within 5 μm of the nucleus (Perinuclear). Perinuclear Ratio was measured as: PR= Perinuclear/ Total.

### Tubule-to-centrosome angle

To ascertain if LYTL tubules are oriented towards the centrosome, the tubule-to-centrosome angle was measured. Briefly, a segmented line was traced following LYTL tubules from their tup to their base, and then from the base to the centrosome (stained with pericentrin). The angle of the segmented line was measured with Fiji (ImageJ, NIH). A tubule oriented towards the centrosome, would have an angle close to 0°.

### Steady/Dynamic tubular event measurement

U2OS treated with NTC or siRILPL1 RNA for 24 h were transfected with 3xflag-LRRK2 and mNeonGreen-JIP4 for 36 h. Cells were then treated with LLOME for 2 h and imaged live. Stacks were taken every 3 s, for a full duration of 4 min. LYTL tubular events were categorized as follows: Steady (when tubules did not undergo significant change in the full duration of the movie) and Dynamic (when tubules retracted completely until their base, or underwent sorting). Steady events recorded for less than 30 s from elongation to the end of the movie were not taken into consideration. Only cells with 5 or more JIP4-positive tubules were imaged.

### Immunostaining

#### Imaging fixed LYTL tubules in fixed cells

Cells were fixed with 4% PFA for 10 mins, permeabilized with PBS/ 0.1% Triton for 10 mins and blocked with 5% Donkey Serum for 1 h at RT. Primary antibodies were diluted in blocking buffer (1% Donkey Serum) and incubated overnight at 4°C. After three 5 min washes with PBS/ 0.1% Triton, secondary fluorescently labeled antibodies were diluted in blocking buffer (1% Donkey Serum) and incubated for 1 hour at RT. Coverslips were washed twice with 1x PBS and an additional two times with dH2O, and mounted with ProLong® Gold antifade reagent (Thermo Fisher).

#### Imaging microtubules in fixed cells

Cells were fixed in methanol for 5 min at -20°C and washed with PBS three times. Cells were permeabilized with PBS/ 0.1% Triton for 10 mins and blocked with 5% Donkey Serum for 1 h at RT. Primary antibodies were diluted in blocking buffer (1% Donkey Serum) and incubated overnight at 4°C. After three 5 min washes with PBS, secondary fluorescently labeled antibodies were diluted in blocking buffer (1% Donkey Serum) and incubated for 1 hour at RT. Coverslips were washed twice with 1x PBS and an additional two times with dH2O, and mounted with ProLong® Gold antifade reagent (Thermo Fisher).

#### Imaging LYTL tubules and microtubules in fixed cells

Cells were fixed with 4% PFA and 0.1% Glutaraldehyde for 10 mins at RT. After 5 washes with PBS, cells were incubated with 0.1% NaBH4 in PBS for 10 mins at RT. After 3 washes, cells were incubated in 0.1% NaBH4 + 5% BSA in PBS for 1 h at 37°C. After 3 washes with PBS, cells were permeabilized with PBS/ 0.1% Triton for 10 mins and blocked with 5% Donkey Serum for 1 h at RT. Primary antibodies were diluted in blocking buffer (1% Donkey Serum) and incubated overnight at 4°C. After three 5 min washes with PBS, secondary fluorescently labeled antibodies were diluted in blocking buffer (1% Donkey Serum) and incubated for 1 hour at RT. Coverslips were washed twice with 1x PBS and an additional two times with dH2O, and mounted with ProLong® Gold antifade reagent (Thermo Fisher).

### RILPL1: RAB10 structural binding model

The X-cap of RILPL1 was created using AlphaFold2 predictive modeling. The model of RAB10 was created based on the coordinates from (Rai et al., 2016) (PDB ID: 5SZJ). Predictive interaction of RLIPL1/RAB10 was created in AlphaFold2 Multimer. ChimeraX software (Pettersen et al., 2021) was used to visualize and render all structures.

### Coimmunoprecipitation

HEK293FT and U2OS cells transfected with 3xflag-LRRK2 and mCherry-RILPL1 plasmids were lysed in IP buffer (20 mM tris-HCl (pH 7.5), 150 mM NaCl, 1 mM EDTA, 0.3% Triton X-100, 10% glycerol, 1× Halt protease and phosphatase inhibitor cocktail) (Thermo Fisher Scientific) for 30 min on ice. Lysates were centrifuged at 4°C for 10 min at 20,000*g*. The supernatant was incubated in RFP-Trap magnetic beads (Chromotek) for 1 hour at 4°C on a rotator. Beads were washed six times with IP wash buffer (20 mM tris-HCl (pH 7.5), 300 mM NaCl, 1 mM EDTA, 0.1% Triton X-100, 10% glycerol) and eluted in 2× loading buffer (Biorad) with β-mercaptoethanol by boiling for 5 min at 95°C.

### SDS PAGE and Western Blotting

Proteins were resolved on 4–20% Criterion TGX pre-cast gels (Biorad) and transferred to membranes by semi-dry trans-Blot Turbo transfer system (Biorad). The membranes were blocked with Intercept Blocking Buffer (LICOR) and then incubated for 1h at RT or overnight at 4°C with the indicated primary antibody. The membranes were washed in TBST (3×5 min) followed by incubation for 1h at RT with fluorescently conjugated goat anti-mouse, rat or rabbit IR Dye 680 or 800 antibodies (LICOR). The blots were washed in TBST (3×5 min) and scanned on an ODYSSEY^®^ CLx (LICOR) and an Azure 500 (Azure). Quantitation of western blots was performed using Image Studio (LICOR) and Azure Spot Pro (Azure)..

### Statistical analysis

Statistical analysis for experiments with two treatment groups used Student’s t-tests with Welch’s correction for unequal variance. F-test was used to assess the equality of variances of t-test. For more than two groups, we used one-way analysis of variance (ANOVA) or two-way ANOVA where there were two factors in the model. Tukey’s post hoc test was used to determine statistical significance for individual comparisons in those cases where the underlying ANOVA was statistically significant and where all groups were compared; Dunnett’s multiple comparison test was used where all groups were compared back to a single control group. Unless otherwise stated, graphed data are presented as means ± SEM or violin plots. Comparisons were considered statistically significant are indicated; *, p< 0.05; **, p< 0.01; ***, p< 0.001; ****, p< 0.0001.

## SUPPLEMENTARY MOVIE LEGENDS

**Supplementary movie 1.** U2OS cell transfected with HaloTag-LRRK2 for 36 h and treated with LLOME. Maximum intensity projection. Cell was imaged for 120 min at 5 min per stack. Movie is played at 7 seconds per frame. Scale bar: 10 µm.

**Supplementary movie 2.** U2OS cells were transfected with 3xflag-LRRK2, mNeonGreen-JIP4 and TUBB5- HaloTag. Subsequently, cells were treated with LLOME (2 h). Cell was imaged for 4 minutes at 3 seconds per stack. Movie is played at 4 seconds per frame. White arrowheads indicate LYTL tubules associated with microtubules and outlined arrowheads show a tubule budding along a microtubule. Scale bar: 2 µm.

**Supplementary movie 3.** U2OS cells were transfected with 3xflag-LRRK2, HaloTag-JIP4 and EMTB- mNeonGreen. Subsequently, cells were treated with LLOME (2 h). Cell was imaged for 4 minutes at 3 seconds per stack. Movie is played at 4 seconds per frame. White arrowheads indicate LYTL sorted material traveling on microtubules. Scale bar: 2 µm.

**Supplementary movie 4.** U2OS cells were transfected with 3xflag-LRRK2, HaloTag-JIP4 and EMTB- mNeonGreen. Subsequently, cells were treated with LLOME (2 h) with (DMSO) or with Nocodazole (10 µM, 30 min). Both cells were imaged for 4 minutes at 3 seconds per stack. Movies are played at 7 seconds per frame. White arrowheads indicate LYTL tubules and outlined arrowheads show sorted material. Scale bar: 2 µm.

**Supplementary movie 5.** U2OS cells were transfected with 3xflag-LRRK2, mNeonGreen-JIP4 and TagRFP- A1aY. Subsequently, cells were treated with LLOME (2 h). Cell was imaged for 105 seconds at 3 seconds per stack. Movies are played at 7 seconds per frame. White arrowheads indicate a sorted tubule moving along a tyrosinated microtubule. Scale bar: 2 µm.

