## Supplementary Figures 1-5 for "Opposing actions of JIP4 and RILPL1 provide antagonistic motor force to dynamically regulate membrane reformation during lysosomal tubulation/sorting driven by LRRK2"

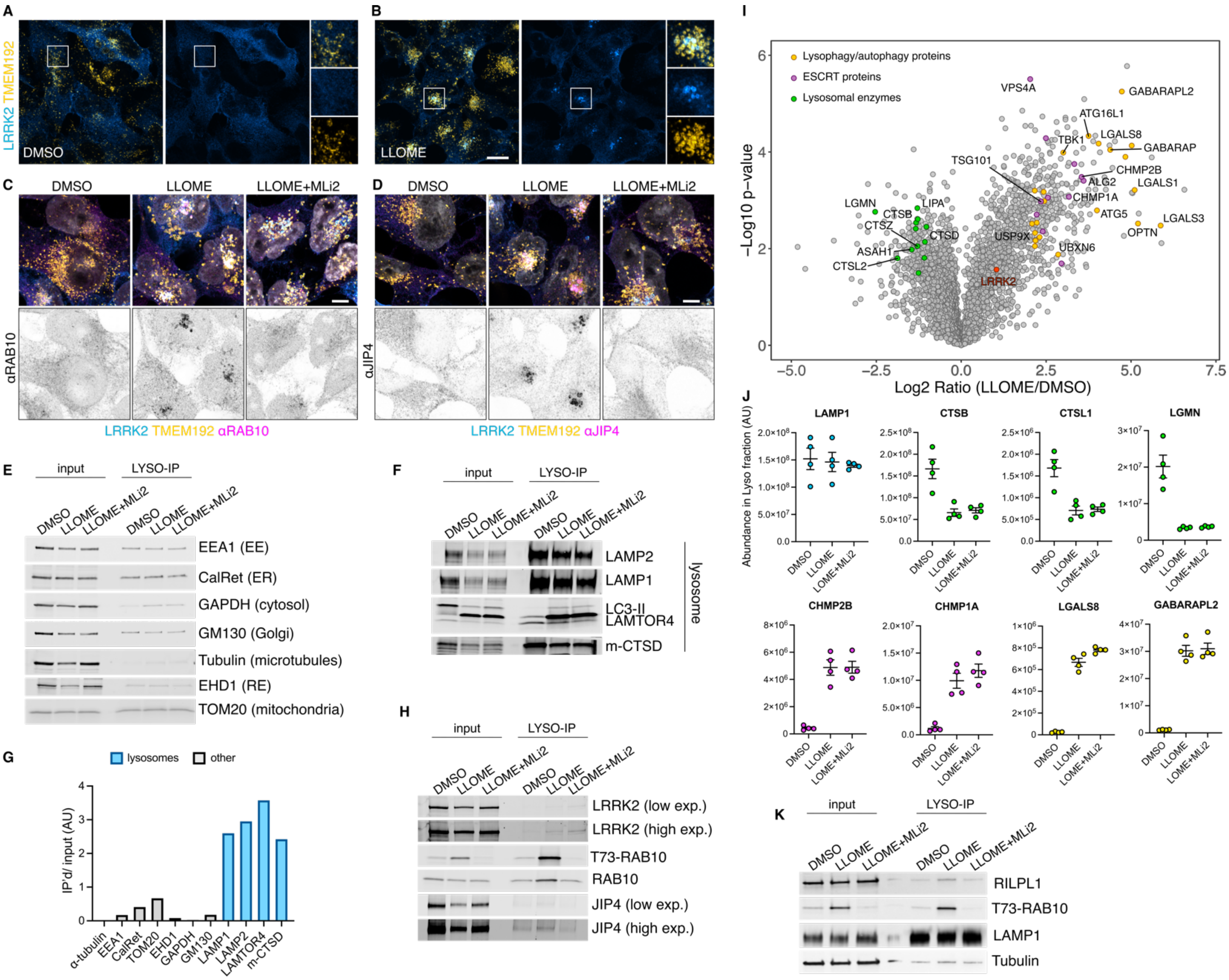


***Supplementary Figure 1. Additional information regarding the LYSO-IP unbiased proteomics screening.*** *(A-B) HEK293T cells stably expressing GFP-LRRK2 and TMEM192-3xHA were seeded for 24 h and treated with DMSO or LLOME for 2 h. Cells were then fixed and stained using an anti HA antibody. (C-D) HEK293T cells stably expressing GFP-LRRK2 and TMEM192-3xHA were seeded for 24 h and treated with DMSO, LLOME (2 h) and LLOME+MLi2. Cells were then fixed and stained for HA and endogenous RAB10 (C), or endogenous JIP4 (D). (E, F, H) HEK293T cells stably expressing GFP-LRRK2 and TMEM192-3xHA were seeded for 24 h and treated with DMSO, LLOME (2 h) and LLOME+MLi2. Cells were then subjected to LYSO-IP to immunoisolate lysosomes. Isolated lysosomes were lysed and immunoblots show the amount of different cellular compartments in the lysosomal fraction, including EE, ER, cytosol, Golgi, RE and mitochondria (E), as well as different luminal and membranous lysosomal proteins (F). (G) Marker quantification confirming enrichment of lysosomal proteins in the lysosomal fraction. (H) Immunoblot from LYSO-IP of players involved in LYTL, including pT73-RAB10, total RAB10, LRRK2 and JIP4. (I) Volcano plot showing the proteins with enhanced recruitment to lysosomes (right side) and the proteins decreased on lysosomes (left side) when cells are treated with LLOME. Data are from 4 independent experiments. (J) Histogram shows protein levels measured by mass spectrometry of individual proteins on lysosomes under DMSO, LLOME and LLOME+MLi2. Data are from 4 independent experiments. (K) Immunoblot from LYSO-IP experiment showing that RILPL1 recruitment to damaged lysosomes is LRRK2 kinase activity dependent.*

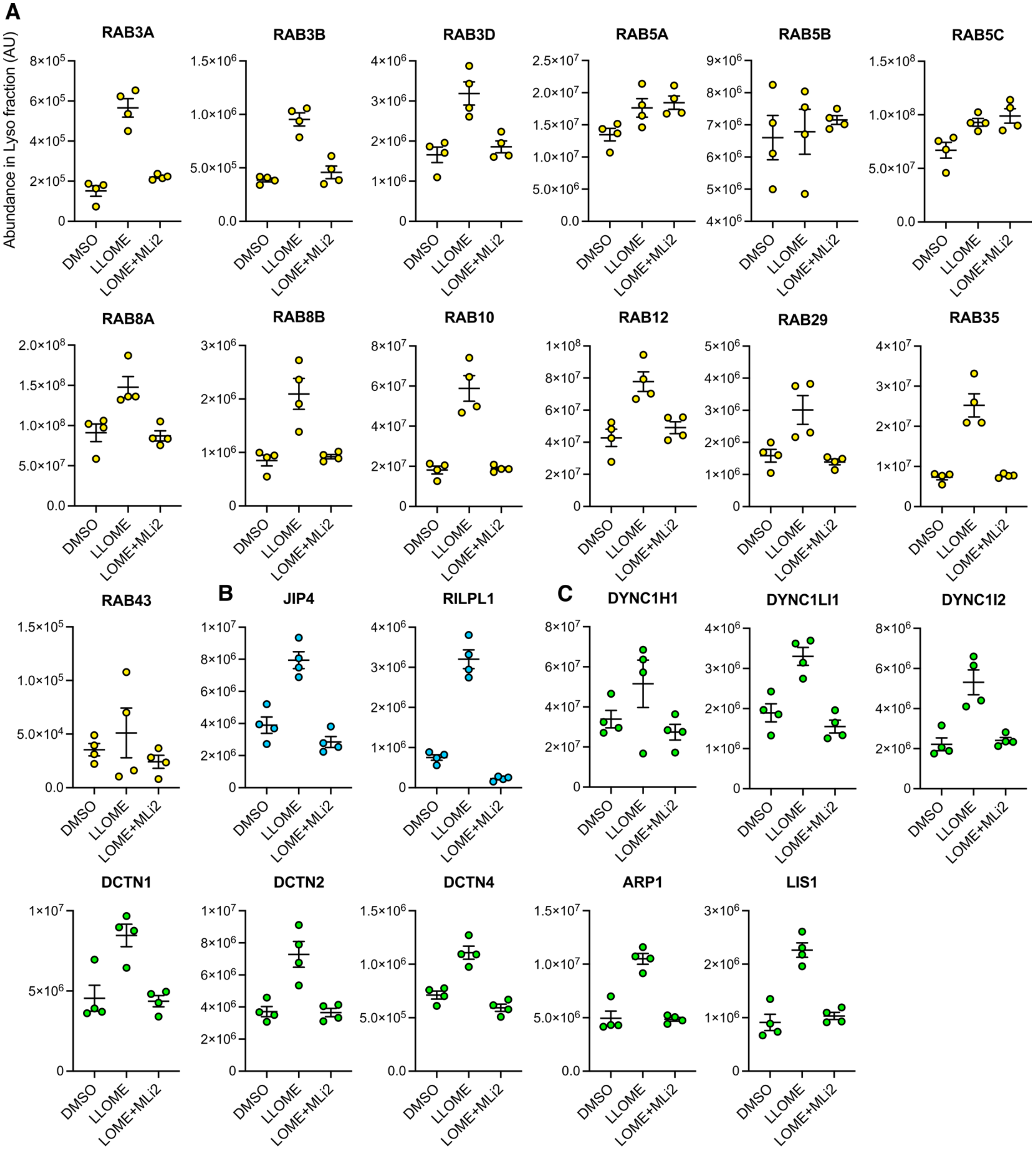


***Supplementary Figure 2. Additional information regarding Figure 1****. Histogram shows protein levels measured by mass spectrometry of individual proteins on lysosomes under DMSO, LLOME and LLOME+MLi2. Data are from 4 independent experiments. (A) RAB substrates, (B) RHD proteins and (C) dynein/dynactin subunits.*

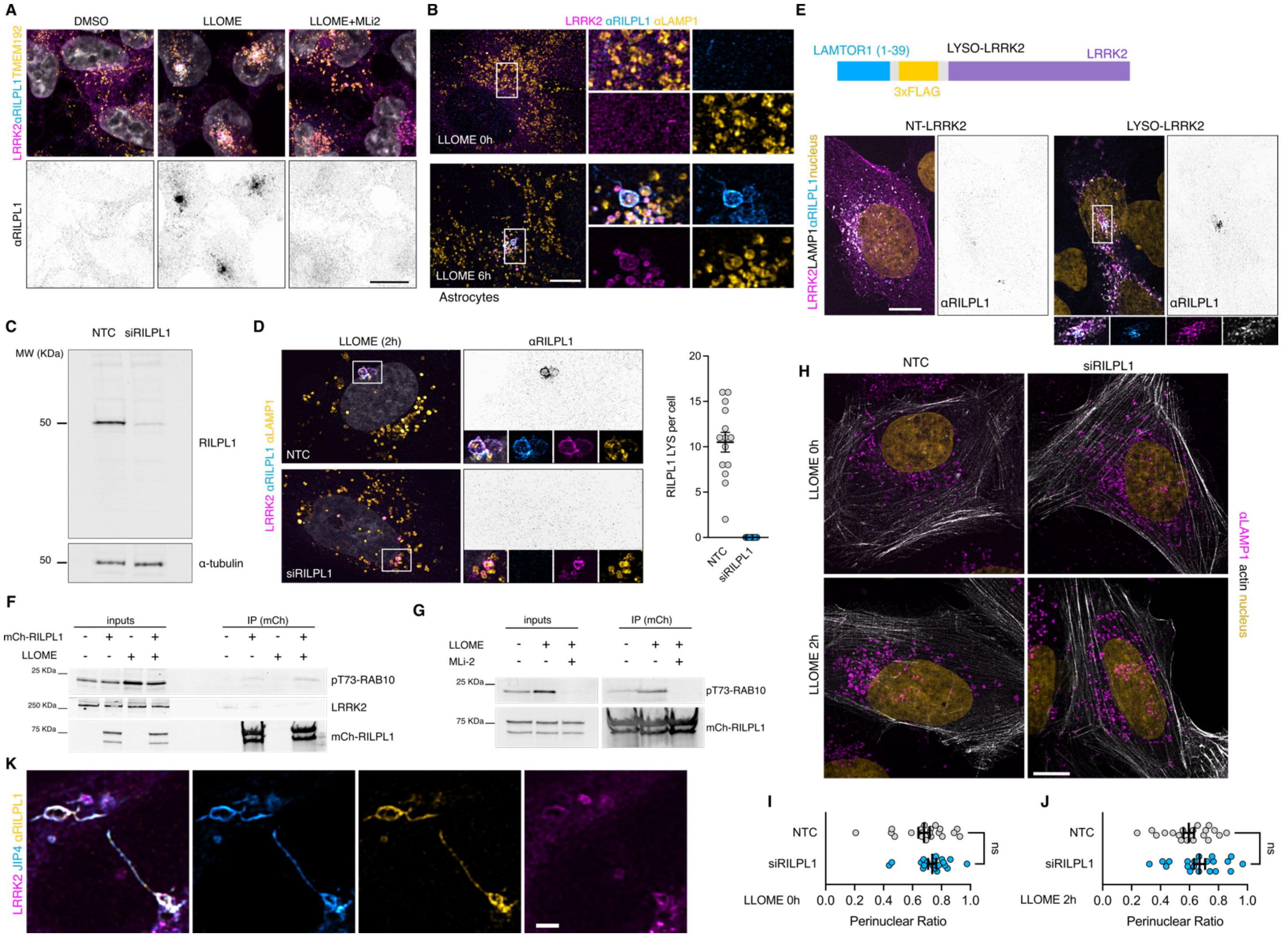


***Supplementary Figure 3. Additional information regarding RILPL1 recruitment to lysosomes by LRRK2****. (A) HEK293T cells stably expressing GFP-LRRK2 and TMEM192-3xHA were seeded for 24 h and treated with DMSO, LLOME (2 h) and LLOME+MLi2. Cells were then fixed and stained for HA and endogenous RILPL1. (B) Mouse primary astrocytes were transfected with 3xflag-LRRK2 for 48 h. Cells were then treated or not with LLOME for 6 h, fixed and stained for endogenous LAMP1 and RILPL1. (C) U2OS cells were treated with a non-targeting control (NTC) or siRILPL1 for 60 h. Cells were then lysed and blotted for endogenous RILPL1. (D) U2OS cells were treated with a non-targeting control (NTC) or siRILPL1 for 24 h. Cells were then transfected with 3xflag-LRRK2 for 36 h, treated with LLOME for 2 h and fixed. Cells were stained for endogenous RILPL1 and LAMP1. Graph shows the number of RILPL1-positive lysosomes per cell in both conditions (n =13-14 cells). (E) U2OS cells were transfected with LAMP1-HaloTag and 3xflag-LRRK2 (NT-LRRK2) or LAMTOR1(1-39)-3xflag-LRRK2 (LYSO-LRRK2) for 36 h. Cells were fixed and stained for endogenous RILPL1. (F-G) U2OS cells were transfected with HaloTag-LRRK2 and mCherry-RILPL1 for 36 h. Cells were then treated or not with LLOME (2 h) (F) or pre-treated with MLi2 (G). Lysates were subjected to immunoprecipitation with anti-RFP antibodies. (H) U2OS cells were treated with a non-targeting control (NTC) or siRILPL1 for 24 h. Cells were then transfected with LifeAct-mNeonGreen for 36 h, treated or not with LLOME for 2 h and fixed. Cells were then stained for endogenous LAMP1. (I-J) Graphs show the perinuclear ratio of lysosomes in the different conditions. Unpaired t-test. Data are Mean ± SEM (n= 20 cells). (K) U2OS cells transfected with HaloTag-LRRK2 and mNeonGreen-JIP4 for 36 h, and treated with LLOME for 2 h before fixing. Cells were then stained for endogenous RILPL1.Scale bar= 10 μm.*

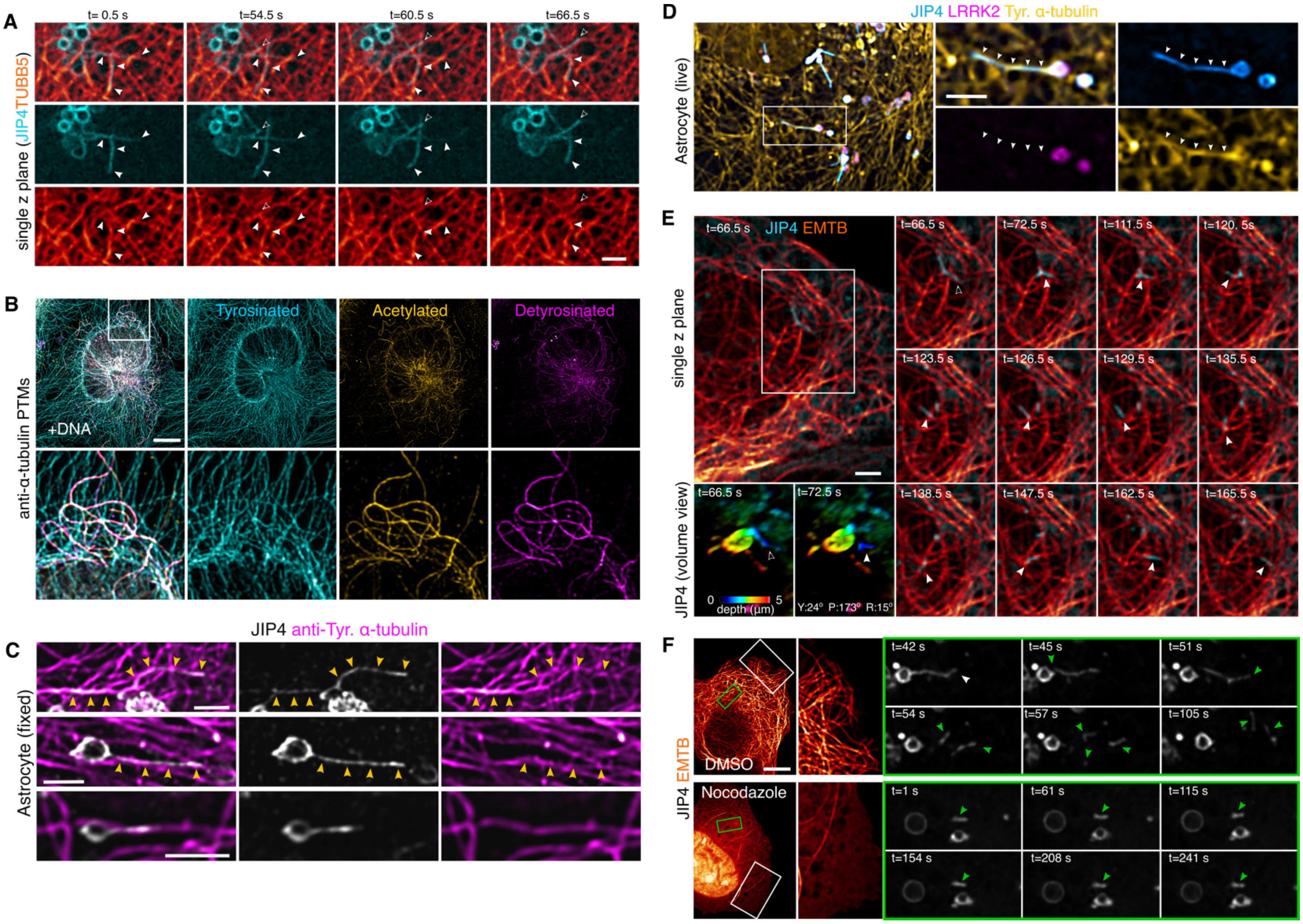


***Supplementary Figure 4. Additional information regarding α-tubulin PTMs contacts with LYTL tubules.*** *(A) U2OS cells were transfected with 3xflag-LRRK2, mNeonGreen-JIP4 and TUBB5-HaloTag. Cells were treated with LLOME (2 h) and observed under a confocal microscope. Time-lapse shows LYTL tubules associated with microtubules (arrowheads) and even a tubule budding alongside a microtubule (outlined arrowhead). (B) U2OS cells were fixed and stained for different α-tubulin PTMs: tyrosinated, acetylated and detyrosinated. (C) Mouse primary astrocytes were transfected with 3xflag-LRRK2 and pDEST53-JIP4 for 48 h. Cells were treated with LLOME (6 h) and fixed. After fixation cells were stained for GFP and tyrosinated α-tubulin. Yellow arrowheads indicate contact between LYTL tubule and tyrosinated microtubules. (D) Mouse primary astrocytes were transfected with HaloTag-LRRK2, mNeonGreen-JIP4 and TagRFP-T-A1aY1 (tyrosinated microtubules) for 48 h. Cells were treated with LLOME (6 h) and imaged live. Arrowheads indicate contact between LYTL tubule and tyrosinated microtubules. (E) U2OS cells were transfected with 3xflag-LRRK2, HaloTag-JIP4 and EMTB-mNeonGreen. Cells were treated with LLOME (2 h) and analyzed under a confocal microscope. Time-lapse shows a sorted tubule moving along microtubules on a single z plane. Volume view with depth code shows the tubule attached to a lysosome (t=66.5 s) and the moment of fission (t=72.5 s). (F) U2OS cells were transfected with 3xflag-LRRK2, HaloTag-JIP4 and EMTB-mNeonGreen. Cells were treated with LLOME (2 h) and with DMSO or Nocodazole (10 µM, 30 min) and analyzed under a confocal microscope. Time-lapse shows the effect of nocodazole on the microtubule network and the dynamics of the LYTL sorted materials. Scale bar (A,C,D,E)= 2 µm; (B,F)= 10 µm.*

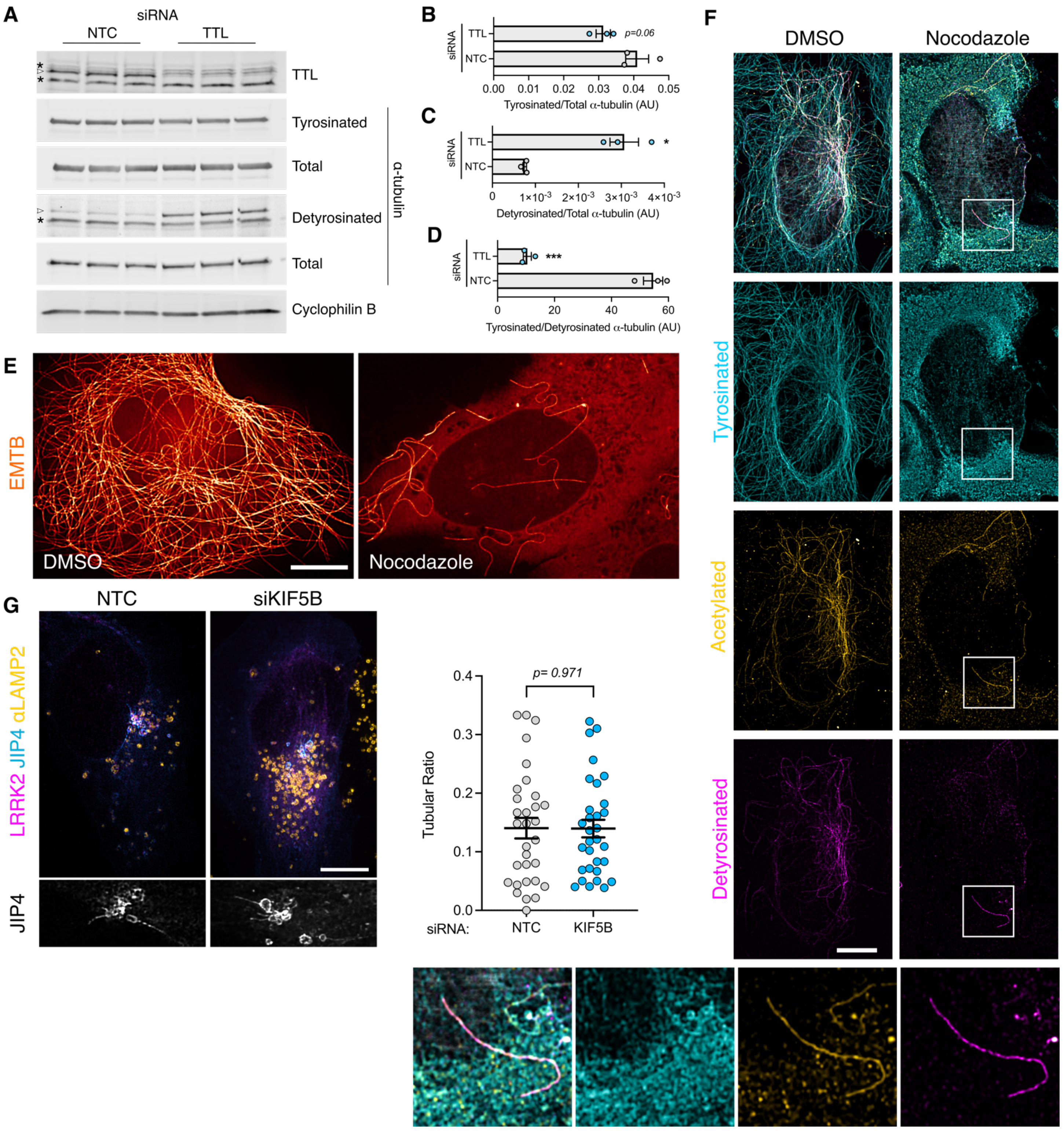


***Supplementary Figure 5. Additional information regarding tyrosinated α-tubulin and LYTL tubule elongation.*** *(A) U2OS cells were transfected with a non-targeting control (NTC) or TTL siRNA for 60 h. Western blot shows tyrosinated α-tubulin, detyrosinated α-tubulin, total α-tubulin and cyclophilin B protein levels. (B) Histogram showing tyrosinated α-tubulin protein levels normalized to total α-tubulin. Unpaired t-test was used (p=0.066, n=3). (C) Histogram showing detyrosinated α-tubulin protein levels normalized to total α-tubulin. Unpaired t-test with Welch's correction was used (p=0.0185, n=3). (D) Histogram showing the normalized tyrosinated/detyrosinated α-tubulin ratio. Unpaired t-test was applied (p=0.0003, n=3). Asterisks signal unspecific bands; real bands shown in arrowhead. (E) U2OS cells were transfected with EMTB-mNeonGreen and treated with DMSO or Nocodazole (10 μM, 30 min) and imaged live under a confocal microscope. (E) U2OS cells were treated with DMSO or Nocodazole (10 μM, 30 min) and fixed. (F) U2OS cells treated with DMSO or Nocodazole were fixed and stained for tyrosinated α-tubulin, detyrosinated α-tubulin and acetylated α-tubulin. (G) U2OS cells were transfected with a non-targeting control (NTC) or KIF5B siRNA for 24 h. Cells were then transfected with HaloTag-LRRK2 and mNeonGreen-JIP4 for 36 h and treated with LLOME for 2 h. Cells were then fixed and stained for endogenous LAMP2. Graph depicts the tubular ratio in both conditions. Unpaired t-test. Data are Mean ± SEM (n= 31 cells). Scale bar= 10 μm.*
